# Phosphorylation of the mRNP component Yra1 couples heat stress to nuclear mRNA export inhibition

**DOI:** 10.64898/2026.06.25.734561

**Authors:** Johanna Franziska Seidler, Jeanette Dalwig, Johannes Graumann, Katja Sträßer

## Abstract

The ability to adapt to changing environmental conditions is essential for cellular survival. A central feature of the eukaryotic stress response is the inhibition of bulk mRNA nuclear export, while stress-induced transcripts are specifically exported. However, the molecular mechanisms that simultaneously inhibit bulk mRNA export while mediating selective export of specific transcripts remain poorly understood. Here, we performed comparative phosphoproteomic analyses of *S. cerevisiae* under different stress conditions. We identified a heat shock-induced increase in phosphorylation within the N-terminal domain of the mRNA export adaptor Yra1. Preventing this phosphorylation significantly reduces nuclear accumulation of poly(A)^+^ RNA during heat stress and concomitantly enhances the export of heat-induced transcripts. Mechanistically, Yra1 phosphorylation appears to weaken its interaction with the export receptor Mex67, thereby contributing to nuclear accumulation of bulk poly(A)^+^ RNA under heat stress. Together, our findings establish Yra1 phosphorylation as a previously unrecognized regulatory mechanism that promotes nuclear mRNA accumulation during heat stress and contributes to selective mRNA export.

## INTRODUCTION

The ability to mount effective adaptive responses to environmental stress is essential for cell survival and varies considerably both between stress conditions and across organisms. A central mechanism underlying such adaptation is the dynamic regulation of gene expression, and among its many layers, mRNA metabolism has emerged as a particularly important target of stress-responsive signaling pathways. In *S. cerevisiae*, several stress conditions—including heat stress, hyperosmotic stress and ethanol treatment—trigger a striking accumulation of poly(A)^+^ within the nucleus, implying a broad inhibition of bulk mRNA export.^1^ Concurrently, stress-specific transcripts are selectively exported to the cytoplasm, where they can be rapidly translated. This selective export-versus-retention dichotomy is not restricted to yeast but has been observed across diverse eukaryotes in response to stress and infection.^2–6^ How cells achieve this remarkable specificity—simultaneously retaining bulk mRNAs while licensing the export of stress-induced transcripts—remains poorly understood. Moreover, whether nuclear mRNA accumulation represents a generic response to cellular stress or is instead restricted to specific stress conditions has not been systematically addressed.

Nuclear mRNA export is an evolutionarily conserved and highly coordinated multi-step process.^7–9^ It begins co-transcriptionally as the nascent transcript emerging from RNA polymerase II is bound by a series of RNA-binding proteins (RBPs) that together assemble into a messenger ribonucleoprotein particle (mRNP) competent for nuclear export.^7,9,10^ A central organizer of nuclear mRNP assembly is the evolutionary conserved TREX complex, which couples transcription elongation to mRNA export.^9^ In *S. cerevisiae*, TREX comprises the pentameric THO complex (Hpr1, Tho2, Thp2, Mft1, Tex1), the DEAD-box helicase Sub2 (human orthologue: UAP56), the RNA annealing protein Yra1 (human orthologue: ALYREF) and the SR-like proteins Gbp2 and Hrb1.^9,11,12^ Deletion of THO complex subunits impairs both transcription elongation and nuclear mRNA export and causes hyperrecombination, reflecting the accumulation of transcription-associated DNA:RNA hybrids.^9,13–15^ Sub2 functions in both pre-mRNA splicing and mRNA export, where its ATPase activity drives nuclear mRNP remodeling^11,16,17^; this activity is stimulated by THO and by Yra1.^18,19^ Yra1 itself harbors conserved N- and C-terminal domains containing UAP56-binding motifs (UBMs) essential for Sub2 interaction^20,21^ and plays a central role in compacting nascent transcripts into export-competent mRNPs.^22,23^ Yra1, the THO component Hpr1, the poly(A)^+^-binding protein Nab2 and the SR-like protein Npl3 function as a mRNA export adaptors: these factors recruit the heterodimeric export receptor Mex67-Mtr2 to the mRNP.^24–27^ Mex67-Mtr2 in turn mediates translocation of the mRNP through the nuclear pore complex (NPC) by engaging FG-nucleoporins lining the central transport channel.^28–31^ On the cytoplasmic face of the NPC, the DEAD-box helicase Dbp5 remodels the exported mRNP by displacing nuclear RBPs, thereby preventing their reimport and rendering the export event irreversible.^8^

Nuclear mRNA export is profoundly altered under stress conditions. While stress-induced transcripts are selectively exported, bulk mRNA accumulates in the nucleus, implying that the two processes are regulated independently. Notably, many canonical mRNA export factors—including THO complex components and export adaptors—appear to be dispensable for the export of heat shock transcripts ^32–34^, suggesting that heat shock mRNA export operates via a streamlined pathway centered on Mex67-Mtr2, Sub2 and nucleoporins. In support of this model, Mex67 is recruited directly to nascent heat shock transcripts through interactions with the heat shock transcription factor Hsf1 and RNA polymerase II, bypassing the need for canonical export adaptors.^33,35^ It has further been proposed that dissociation of export adaptors from the mRNA export machinery during stress contributes to the nuclear retention of bulk mRNAs. Nevertheless, several lines of evidence complicate this picture: Yra1 associates with the heat shock transcript *SSA4*^32^, and deletion or mutation of export adaptors or THO complex subunits causes nuclear *SSA4* foci ^32,36^, arguing that these factors do contribute to heat shock mRNA export under certain conditions. A key regulatory role for the MAP kinase Slt2, involved in the cell wall integrity (CWI) pathway, in heat stress-induced mRNA export inhibition was established by the finding that *Δslt2* cells fail to accumulate nuclear poly(A)^+^ RNA during heat stress.^37^ Slt2 phosphorylates the export adaptor Nab2 at T178 and S180; however, preventing phosphorylation at these residues does not abolish nuclear poly(A)^+^ RNA accumulation^37^, indicating that additional Slt2 substrates mediate this response. The identity of these substrates and the precise molecular mechanism by which Slt2 inhibits bulk mRNA export during heat stress have remained unknown.

Here, we systematically address these questions. We demonstrate that nuclear poly(A)^+^ RNA accumulation is a highly stress-specific phenomenon and map the phosphoproteomic landscape of mRNA export factors under heat and hyperosmotic stress. Through this analysis, we identify phosphorylation of the export adaptor Yra1 at S8 – within its conserved N-terminal UBM – as a critical and Slt2-dependent regulatory event. Preventing S8 phosphorylation reduces nuclear poly(A)^+^ RNA accumulation during heat stress, enhances the Yra1-Mex67 interaction and increases Mex67 association with nuclear mRNPs. Unexpectedly, Yra1-S8 phosphorylation is also required for the efficient expression of heat stress-induced transcripts and for efficient splicing. These findings reveal that Yra1-S8 phosphorylation serves a dual function during heat stress: promoting the expression of stress-induced transcripts while simultaneously restricting their nuclear export. We thus identify phosphorylation of the Yra1 N-terminal UBM as a novel regulatory mechanism that coordinates nuclear mRNP assembly and selective mRNA export in response to heat stress.

## RESULTS

### Nuclear Accumulation of mRNA Is Stress-Specific

It is well established that nuclear export of bulk mRNA is inhibited in response to heat shock, hyperosmotic stress, ethanol stress and glucose withdrawal.^1,38,39^ To determine whether this nuclear mRNA accumulation represents a universal stress response, we exposed cells to a broader range of stress conditions and analyzed poly(A)⁺ RNA localization by oligo(dT) fluorescence *in situ* hybridization (FISH) (Figures 1A, 1B, and S1A). In untreated control cells, poly(A)⁺ RNA is uniformly distributed throughout the cytoplasm, with little to no signal detected in the vacuole or nucleus (Figure 1A). Consistent with previous reports, heat, hyperosmotic and ethanol stress each induce robust nuclear mRNA accumulation, as reflected by a pronounced increase in the nuclear-to-cytoplasmic poly(A)⁺ RNA signal ratio.^1^ Hypoosmotic stress, glucose starvation and nitrogen starvation similarly cause nuclear mRNA accumulation (Figure 1A). By contrast, cells treated with rapamycin, arsenite or DTT export mRNA at least as efficiently as wild-type (wt) cells, if not slightly more so (Figure 1A).

**Figure 1.**
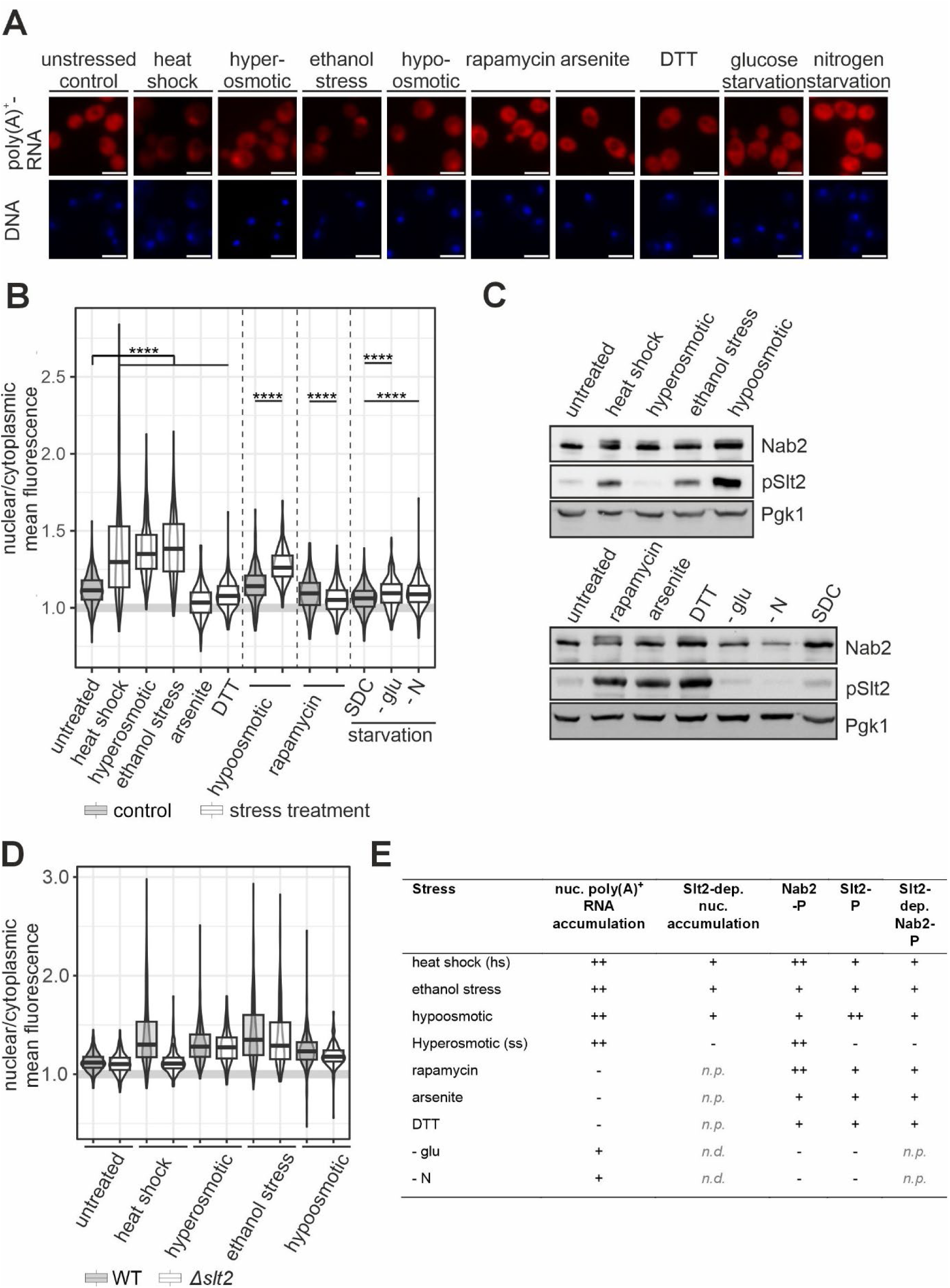
Nuclear accumulation of mRNA is stress-specific. (A) mRNA accumulates in the nucleus under heat, hyperosmotic, ethanol and hypoosmotic stress as well as glucose and nitrogen starvation, but not in response to the other stress conditions. Representative microscopy images of an oligo(dT)-Cy3 FISH to determine the localization of poly(A)^+^ RNA (red). DNA was stained with DAPI (blue). Scale bars: 5 µm. Stress conditions are listed in Table S1. (B) Quantification of the nuclear-to-cytoplasmic ratio of the mean Cy3 fluorescence intensities of the experiments shown in (A). A minimum of 59 cells were quantified per condition and experiment (n ≥ 3). Boxplots display the first and third quartiles (lower and upper hinges), with whiskers extending to the most extreme data points within 1.5 times the interquartile range (IQR). The median is indicated by a horizontal line. Experiments conducted with different controls are separated by dashed lines; representative microscopy images of the controls for hypoosmotic rapamycin treatment and nutrient starvation are shown in Figure S1. Statistical significance: ⃰⃰ p < 0.05, ⃰⃰ ⃰⃰ p < 0.01, ⃰⃰ ⃰⃰ ⃰⃰ p < 0.001, ⃰⃰ ⃰⃰ ⃰ ⃰⃰ p < 0.0001 by Welch’s *t*-test. (C) Phosphorylation of the mRNP component Nab2 and its kinase Slt2 is stress-specific. Nab2 and Slt2 phosphorylation under the indicated stress conditions was assessed by Western blotting. Nab2 phosphorylation was detected by a mobility shift to a higher molecular weight, while phosphorylated Slt2 was specifically recognized by a phospho-specific antibody. (D) Nuclear mRNA accumulation under heat, ethanol and hypoosmotic stress depends on Slt2. Quantification of the nuclear-to-cytoplasmic ratio of mRNA localization in wt (grey) and *Δslt2* (white) cells under control conditions (untreated) and stress conditions that cause a strong impairment of nuclear mRNA export (as shown in Figure 1 A & B). A minimum of 37 cells were quantified per condition in each experiment (n = 3). Statistical significance: *p < 0.05, ⃰⃰ ⃰⃰ p < 0.01, ⃰⃰ ⃰ ⃰⃰ p < 0.001, Representative microscopy images are shown in Figure S1 C. ⃰ ⃰⃰ ⃰⃰ p < 0.0001 by Welch’s *t*-test. (E) Summary of nuclear mRNA accumulation and Nab2 and Slt2 phosphorylation across the tested stress conditions.

Under heat stress, the kinase Slt2 is required for nuclear poly(A)⁺ RNA accumulation, and the mRNP component Nab2 is phosphorylated by Slt2.^37^ We therefore examined both Nab2 phosphorylation and Slt2 activity, as determined by its phosphorylation, across all stress conditions by immunoblotting (Figure 1C). With the exception of hyperosmotic stress, Nab2 phosphorylation correlates with Slt2 activation. However, the magnitude of Nab2 phosphorylation varied considerably across conditions: heat shock and rapamycin treatment induced strong Nab2 phosphorylation, whereas hyperosmotic, hypoosmotic, ethanol, arsenite and DTT stress produce only weak phosphorylation (Figure 1C). Neither glucose nor nitrogen starvation triggers detectable Nab2 phosphorylation or Slt2 activation.

To assess whether Slt2 is required for nuclear mRNA accumulation under conditions that elicit strong nuclear mRNA accumulation as well as Nab2 phosphorylation, we compared nuclear mRNA accumulation and Nab2 phosphorylation in wt and *Δslt2* cells subjected to heat, hyperosmotic, ethanol and hypoosmotic stress (Figure 1D; Figures S1B and S1C). Deletion of *SLT2* reduces the nuclear-to-cytoplasmic poly(A)⁺ RNA signal under heat stress to near-basal levels, and partially attenuated nuclear mRNA accumulation under ethanol and hypoosmotic stress yet has no significant effect on the nuclear poly(A)⁺ RNA retention triggered by hyperosmotic stress. At the biochemical level, *SLT2* deletion abolishes Nab2 phosphorylation under heat, hypoosmotic, rapamycin, arsenite and DTT stress, reduced it under ethanol stress, but left Nab2 phosphorylation under hyperosmotic stress essentially unchanged (Figure S1B).

Taken together, these findings demonstrate that nuclear mRNA accumulation is not a universal stress response but is instead highly stress-specific (Figure 1E). Strikingly, Slt2 activation and Nab2 phosphorylation are neither sufficient nor necessary for nuclear mRNA accumulation. On one hand, rapamycin, arsenite and DTT treatment activate Slt2 and promote Nab2 phosphorylation without causing any detectable nuclear poly(A)⁺ RNA retention (Figure 1E). On the other hand, nutrient starvation drives nuclear mRNA accumulation independently of both Slt2 activity and Nab2 phosphorylation (Figure 1E). Furthermore, while hyperosmotic stress induces nuclear mRNA accumulation accompanied by Nab2 phosphorylation, neither depends on Slt2 (Figure 1E). Furthermore, both the nuclear mRNA accumulation and Nab2 phosphorylation under heat, ethanol and hypoosmotic stress are Slt2-dependent (Figure 1E). These observations reveal an unexpected complexity in the regulation of nuclear mRNA export during stress, suggesting that multiple, mechanistically distinct pathways converge on nuclear mRNA retention in a stress-specific manner.

### Proteins Involved in mRNA Export Undergo Stress-Specific Changes in Phosphorylation

To gain mechanistic insight into stress-induced mRNA export inhibition, we selected heat and hyperosmotic stress as representative conditions, as both elicit robust nuclear poly(A)⁺ RNA accumulation through apparently distinct mechanisms—Slt2-dependent and Slt2-independent, respectively. Since phosphorylation-dependent signaling likely underlies both pathways, we performed quantitative phosphoproteomic analyses of wt cells under control conditions (30°C), heat stress (42°C) and hyperosmotic stress (Figure 2A). To further delineate the contribution of Slt2 to heat stress-induced phosphorylation changes, we extended this analysis to *Δslt2* cells at 30°C and 42°C (Figure 2A). Many sites on proteins are differentially phosphorylated under heat or hyperosmotic stress compared to untreated cells (Figure 2B). Strikingly, the phosphorylation pattern of proteins annotated to the mRNA export pathway (GO:0006406) is clearly distinct between the two stress conditions, reinforcing the conclusion that heat stress and hyperosmotic stress engage mechanistically divergent pathways to drive nuclear poly(A)⁺ RNA accumulation (Figure 2B).

**Figure 2.**
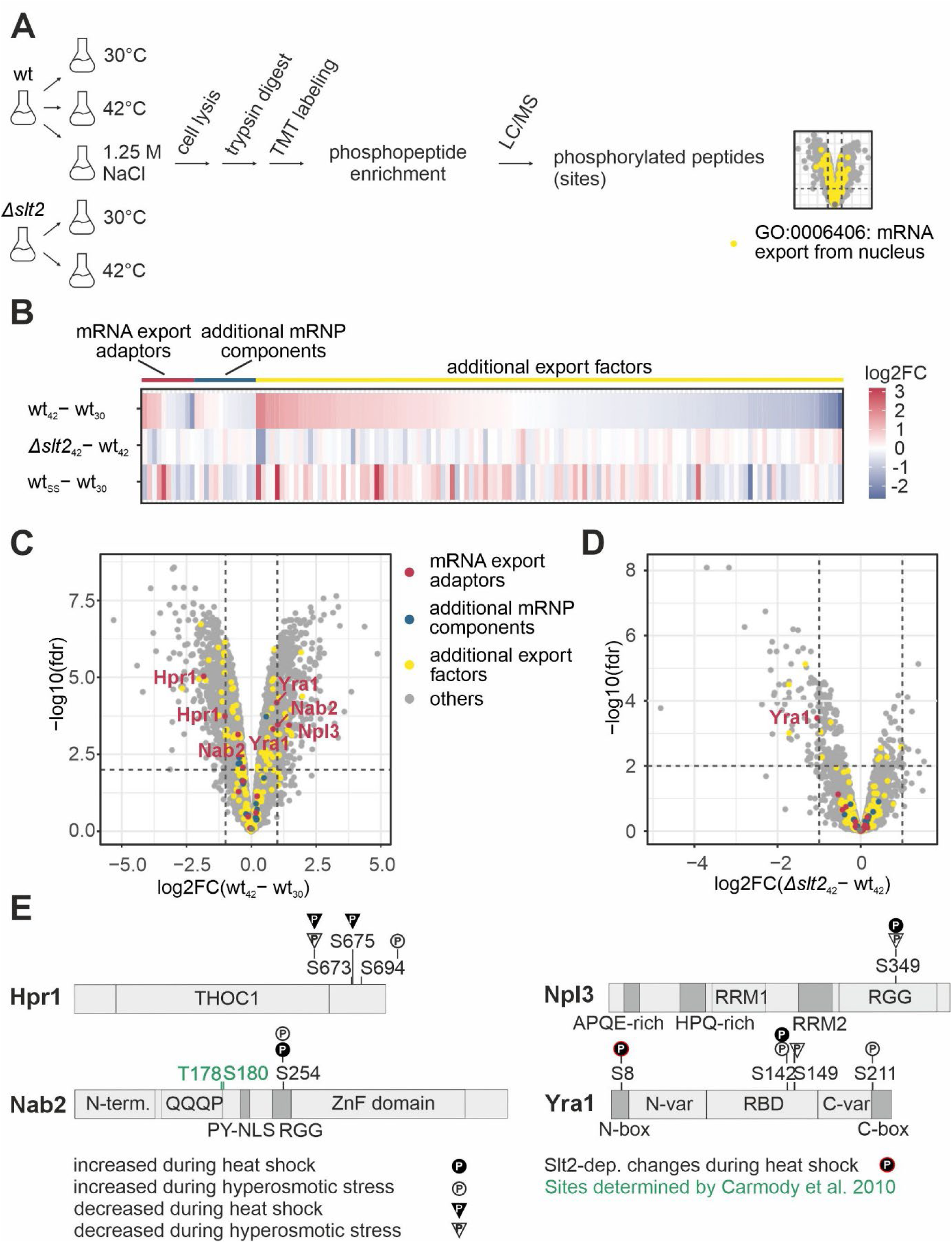
Nuclear mRNA export factors are dynamically phosphorylated in response to heat shock and hyperosmotic stress. (A) Experimental workflow of the phosphoproteome profiling under heat and salt stress conditions. (B) Phosphorylation of nuclear mRNA export factors, including mRNP components, is differentially up- and downregulated in response to heat stress or hyperosmotic stress. Phosphorylation by the kinase Slt2 was assessed in a *Δslt2* strain. Heat map displaying the log_2_-fold changes (log_2_FC) of individual phosphorylation sites in response to heat stress (42°C), hyperosmotic (salt) stress (ss) and in a Slt2 deletion strain grown at 42°C (*Δslt2*_42_). Only sites with a significant p-value (FDR < 0.05) are shown. Multiple bars may correspond to one protein containing several phosphorylation sites. Colored bars above the heat map indicate the classification of each protein: the mRNA export adaptors Nab2, Npl3, Yra1 and Hpr1 in red, additional nuclear mRNP components including additional TREX components, Tho1 and the nuclear cap-binding complex (CBC) in petrol and additional export factors annotated under GO:0006406 (2021) in yellow. (C) Volcano plot showing phosphorylation site changes in response to heat stress in wt cells. Volcano plot depicting the log_2_-fold changes (log_2_FC) in phosphorylation sites in wt cells exposed to 42°C compared to 30°C, plotted against the negative log_10_ of the false discovery rate (FDR). For corresponding data under hyperosmotic stress, see Figure S1. (D) Phosphorylation of specific sites under heat shock is Slt2-dependent, while others are not. Volcano plot as in panel C, comparing phosphorylation site changes between wt and *Δslt2* cells at 42°C. (E) Schematic representation of phosphorylation changes in the four *S. cerevisiae* mRNA export adaptors during heat stress. Increased and reduced phosphorylation during heat stress is indicated by a black circle and triangle, respectively. Increased and decreased phosphorylation during hyperosmotic stress are indicated by a white circle and triangle, respectively. Slt2-dependent phosphorylation during heat stress is indicated by a red border. Fold changes of the respective phosphorylation sites are given in Table S2. Sited identified in a previous study are indicated in green.^37^

Volcano plot analysis of the heat stress phosphoproteome reveals both increased and decreased detection of phosphorylation sites at multiple residues within mRNA export factors (GO:0006406), many of which proved to be Slt2-dependent (Figures 2C and 2D). Notably, all four *S. cerevisiae* mRNA export adaptors display differential phosphorylation at 42°C or under hyperosmotic stress, with nine sites reaching statistical significance (Figure 2E). Specifically, Hpr1 shows reduced phosphorylation at two residues and increased phosphorylation at one residue; Nab2 exhibits increased phosphorylation at one site which does not correspond to the previously identified Slt2 target sites;^37^ Npl3 displays increased phosphorylation at one residue under heat stress and decreased phosphorylation at the same residue under hyperosmotic shock; and Yra1 shows increased phosphorylation at S8 and S142 during heat stress, and at S142 and S211 during hyperosmotic stress, alongside decreased phosphorylation at S149 under hyperosmotic shock. Importantly, of all the differentially phosphorylated sites identified across the four export adaptors, only Yra1-S8 phosphorylation is significantly reduced in *Δslt2* cells at 42°C (Figure 2E), identifying it as the sole Slt2-dependent phosphorylation event among the export adaptors.

In summary, our phosphoproteomic analyses uncover widespread stress-specific remodeling of the phosphorylation landscape of nuclear mRNA export factors, consistent with a broad regulatory reorganization of the export machinery during stress. Given that Yra1-S8 is the only phosphorylation site that decreases in *Δslt2* cells, we focused on this residue as a prime candidate for mediating Slt2-dependent mRNA export inhibition during heat stress.

### Phosphorylation of yra1-S8 is necessary for nuclear mRNA accumulation during heat stress

Our phosphoproteomic analysis identified increased phosphorylation of Yra1 at S8 and, to a lesser extent, at S142 during heat stress. To assess the functional relevance of these sites, we generated a panel of Yra1 mutants carrying either phospho-blocking (serine-to-alanine) or phosphomimetic (serine-to-aspartate) substitutions and examined poly(A)⁺ RNA localization by oligo(dT) FISH (Figures 3A–D and S2).

**Figure 3.**
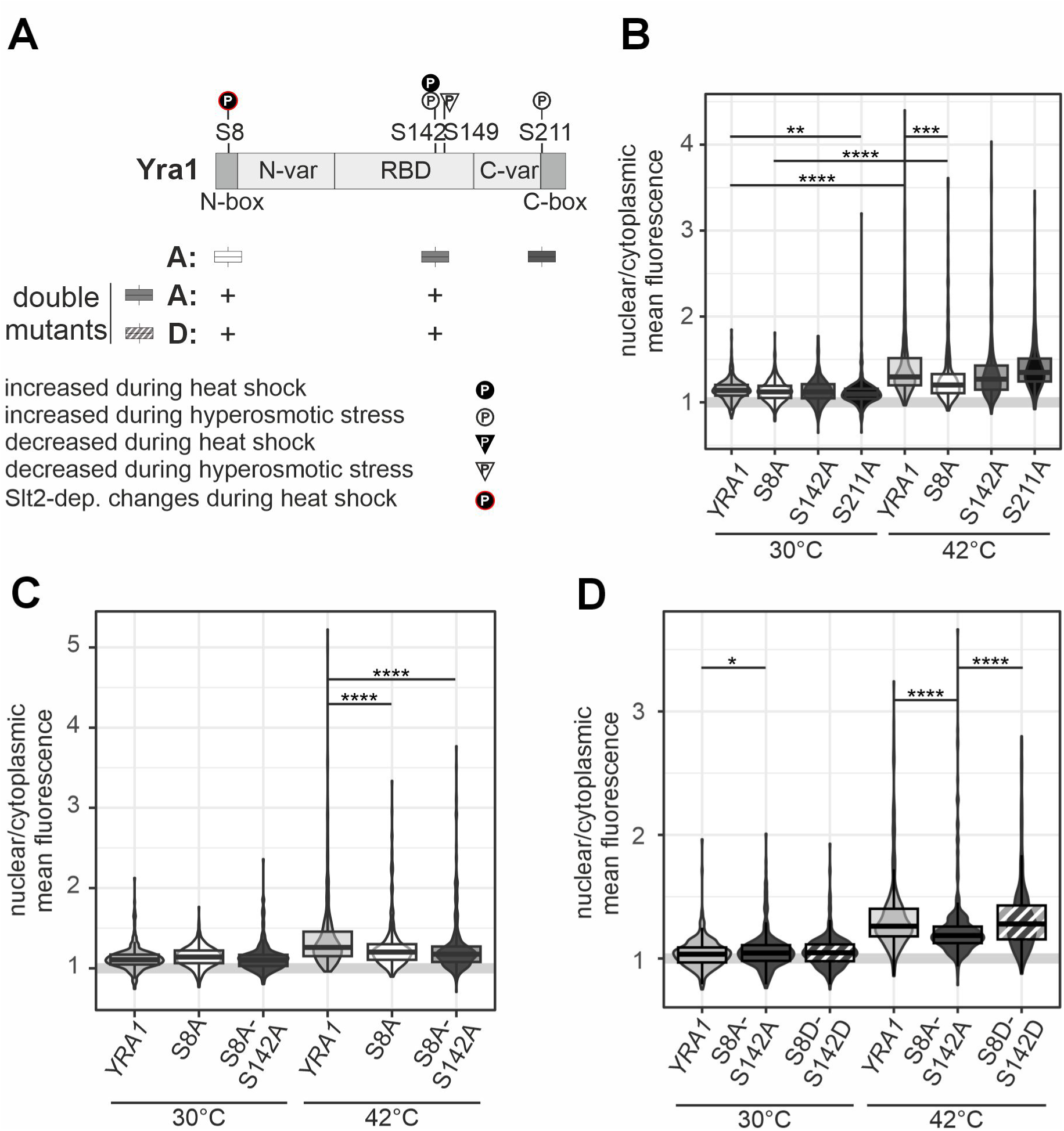
Phosphorylation of the mRNA export adaptor Yra1 at serine 8 in is critical for efficient heat stress-induced nuclear poly(A)^+^ RNA accumulation. (A) Schematic of the Yra1 domain architecture depicting the mutants analyzed in this study. Mutations are denoted using the one-letter amino acid code with A and D indicating the mutation of serine 8, 142 or 211 to alanine and aspartic acid, respectively. Different shades of grey correspond to each mutant and are used in panels B to D. (B-D) Mutation of serine 8 (S8) to alanine (*yra1-S8A*), which prevents phosphorylation at this site, reduces the nuclear accumulation of poly(A)^+^ RNA in response to heat shock. The *yra1-S8A* and the *yra1-S8A-S142A* mutants exhibit decreased nuclear accumulation of poly(A)^+^ RNA under heat stress (B and C). In contrast, the *yra1-S142A* and *yra1-S211A* mutants show nuclear poly(A)^+^ RNA levels comparable to wt cells (B). The phosphomimetic *yra1-S8D-S142D* mutant restores the nuclear accumulation of poly(A)^+^ RNA to wt levels (D). Quantifications of the nuclear-to-cytoplasmic ratio of poly(A)^+^ mRNA localization determined by FISH as in Figure 1. For each experiment, a minimum of 65 cells per sample were analyzed (n ≥ 3). Statistical significance: ⃰ p < 0.05, ⃰⃰ ⃰⃰ p < 0.01, ⃰⃰ ⃰⃰ ⃰⃰ p < 0.001, ⃰⃰ ⃰⃰ ⃰ ⃰⃰ p < 0.0001 by Welch’s t test. Representative microscopy images are shown in Figure S2.

Among the phospho-blocking mutants, only *yra1-S8A* results in a significant reduction in the nuclear-to-cytoplasmic poly(A)⁺ RNA ratio under heat stress (Figure 3B and S2A), indicating that phosphorylation at S8 is required for partial inhibition of nuclear mRNA export. By contrast, the *yra1-S142A* and *yra1-S211A* single mutants have no discernible effect on nuclear poly(A)⁺ RNA accumulation (Figure 3B), arguing against a major role for these sites in the heat stress response. Consistent with S8 being the functionally dominant residue, the *yra1-S8A-S142A* double mutant does not further reduce nuclear poly(A)⁺ RNA accumulation beyond that observed in the *yra1-S8A* single mutant (Figure 3C and S2B). Notably, the phosphomimetic *yra1-S8D-S142D* mutant shows no significant difference from the wt control under heat stress (42°C) conditions, suggesting that the aspartate substitution faithfully recapitulates the effect of phosphorylation of S8. Of note, nuclear mRNA export is unaltered in *yra1-S8D-S142D* cells, indicating that the negative charge at S8 and S142 is insufficient to alter nuclear mRNA export and that additional molecular changes are required (Figure 3D and S2C).

Taken together, these results identify Yra1-S8 phosphorylation as a key regulatory event in the heat stress-induced nuclear mRNA export inhibition. The partial but significant rescue of nuclear mRNA export conferred by the *yra1-S8A* mutation places Yra1-S8 downstream of Slt2 as an important effector of stress-induced nuclear poly(A)⁺ RNA retention.

### Yra1-S8 phosphorylation promotes dissociation of Mex67 from the nuclear mRNP

To determine how Yra1 phosphorylation impacts nuclear mRNA export, we investigated whether phosphorylation at S8 at 42°C affects the composition of nuclear mRNPs. To this end, we purified nuclear mRNPs by purification of the nuclear cap-binding protein Cbp20 together with its associated mRNA and proteins from cells expressing either the phospho-blocking *yra1-S8A-S142A-3xHA* mutant at 42°C (Figures 4A and 4B) or the phosphomimetic *yra1-S8D-S142D-3xHA* mutant at 30°C (Figure S3D).

**Figure 4.**
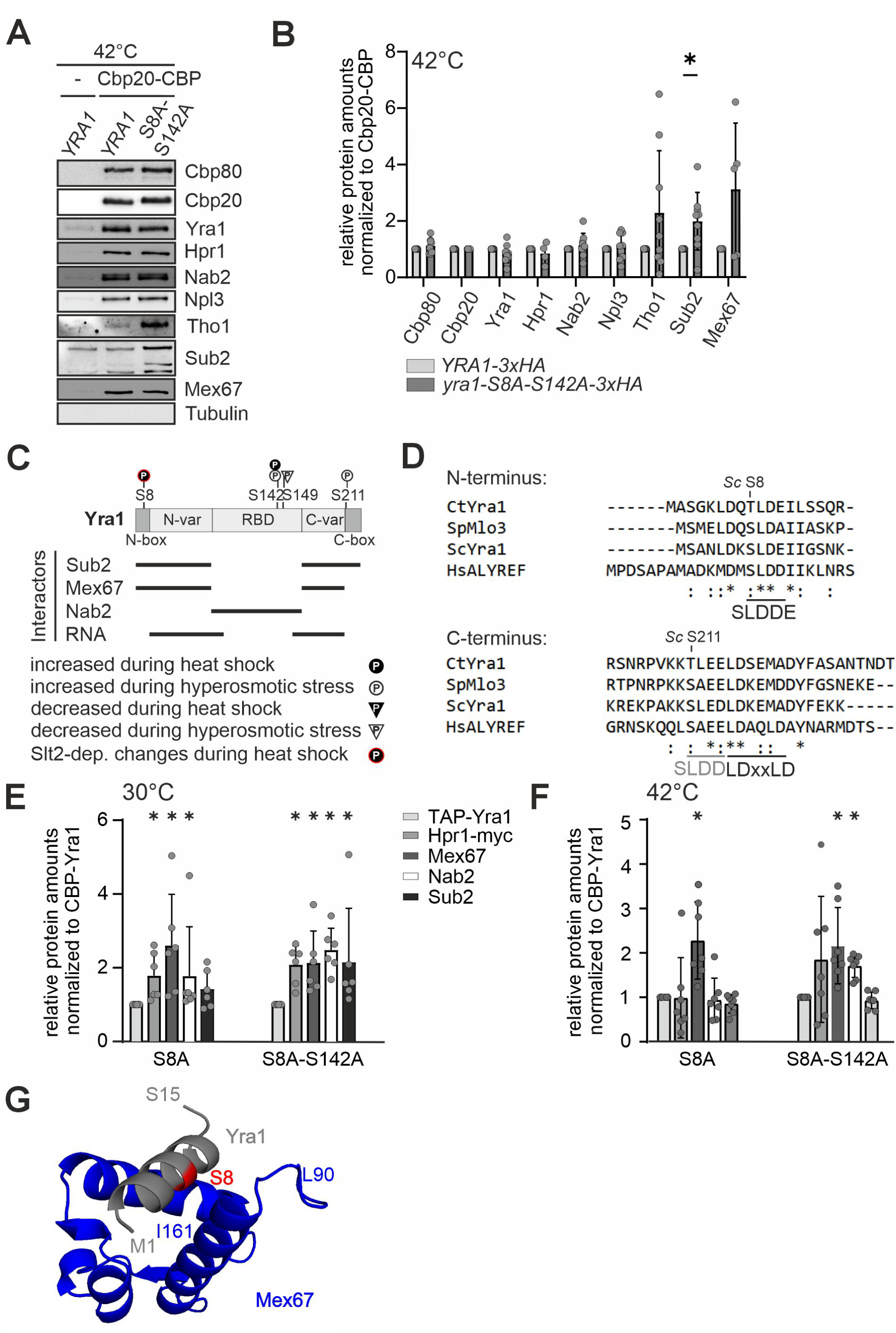
Nuclear mRNPs contain more Mex67 during heat stress in *yra1-S8A-S142A* cells. (A and B) Nuclear mRNPs contain more Mex67 in *yra1-S8A-S142A* cells during heat stress. (A) Representative Western blot of native purifications of nuclear mRNPs via Cbp20-TAP from *YRA1-3xHA* and *yra1-S8A-S142A-3xHA* cells after heat stress (42°C). Co-purifying mRNP components are detected using the indicated antibodies; tubulin served as a negative control. (B) Quantification of the experiment shown in (A). Signal intensities are normalized to the signal intensity of purified Cbp20-CBP and calculated relative to the *YRA1-3xHA* control for each replicate individually. Western blots to determine the total amount of each protein in whole cell lysates are shown in Figure S4. Statistical significance: ⃰ p < 0.05 by Wilcoxon matched-pairs signed rank t-test. (C) Schematic of the Yra1 domain architecture highlighting the phosphorylation sites identified in this study as well as known protein- and RNA-interaction motifs. The positions of the phosphorylation sites including their Slt2 dependency are indicated. The interaction domains of Yra1 with Sub2, Mex67, Nab2 and RNA are indicated by black lines below. (D) Sequence alignment of the N- and C-terminal regions of Yra1 from *S. cerevisiae* and its orthologs in *C. thermophilum* (Yra1), *S. pombe* (Mlo3) and *H. sapience* (ALYREF). The S8 phosphorylation site is located within the conserved UAP56-binding motif (UBM), as identified by Gromadzka et al. (2016)^21^ (indicated with black letters), the S211 residue is located in a putative analogous motif indicated with grey letters. (E) The interaction of Yra1 with Mex67 is enhanced in *yra1-S8A* cells at 30°C. Quantification of the eluates from tandem affinity purifications (TAP) of TAP-Yra1, -yra1-S8A and -yra1-S8A-S142A purifications from cells grown at 30°C. Co-purifying proteins were analyzed by Western blot. Protein amounts were normalized over the amount of Yra1 purified and to the TAP-Yra1 control for comparison for each replicate individually. For representative Western blots and total lysate controls, see Figure S4. Statistical significance: ⃰ p < 0.05by Wilcoxon matched-pairs signed rank t-test. (F) The enhanced interaction of Yra1-S8A with Mex67 persists under heat stress (42°C). Quantification of the eluates from TAP-yra1-S8A and -yra1-S8A-S142A purifications from cells grown at 42°C. Protein amounts were normalized over the amount of Yra1 purified and to the TAP-Yra1 control for comparison. For representative Western blots and total samples see Figure S4. Statistical significance: ⃰ p < 0.05, ⃰⃰ ⃰⃰ p < 0.01, ⃰⃰ ⃰⃰ p < 0.001, ⃰ ⃰⃰ ⃰ ⃰⃰ p < 0.0001 by multiple unpaired t-test. (G) S8 of Yra1 is located in the interaction domain of Yra1 with Mex67. Alphafold model^40^ of the interaction between Ms1 to S15 of Yra1 (gray) with Mex67 L90 to I161 (blue). A model of full-length Yra1 with Mex67-Mtr2 is shown in Figure S4.

Strikingly, preventing Yra1 phosphorylation during heat stress (*yra1-S8A-S142A*) results in a increased association of Mex67 with nuclear mRNPs (Figures 4A and 4B), indicating that phosphorylation of Yra1 at S8 is required for the dissociation of Mex67 from the nuclear mRNP during heat stress. A significant increase in Sub2 association is also observed under these conditions (Figures 4A and 4B). We note that combined phospho-blocking at both S8 and S142 modestly reduces Yra1-S8A-S142A protein levels during heat stress (Figure S3A–C). Conversely, the phosphomimetic *yra1-S8D-S142D* mutant shows a reduction of Yra1 itself in nuclear mRNPs, but no decrease in Mex67 association at 30°C (Figure S3D).

Taken together, these findings demonstrate that phosphorylation of Yra1 at S8 during heat stress likely causes dissociation Mex67 from the nuclear mRNP, providing a mechanistic basis for the Slt2-dependent inhibition of nuclear mRNA export.

### Phosphorylation of Yra1-S8 Promotes Dissociation of Mex67 from Yra1

The Yra1-binding partners Mex67 and Sub2 interact with both the N- and C-terminus of Yra1. Notably, the heat stress-specific phosphorylation site S8 and the hyperosmotic stress-specific site S211 are located in the N- and C-terminus, respectively (Figure 4C). Alignment of multiple UAP56/Sub2 homologs revealed four conserved UAP56-binding motifs (UBMs): SLDD/E, LDxxLD, HDxR and WxHD.^21^ In human ALYREF (the Yra1 orthologue), three such UBMs were identified: an SLDD motif in the N-box, a WxHD motif in the central region and an LDxxLD motif in the C-box. Sequence alignment of *S. cerevisiae* and *Chaetomium thermophilum* Yra1 with *Schizosaccharomyces pombe* Mlo3 and human ALYREF reveals conservation of these motifs in both the N- and C-box (Figure 4D). Strikingly, the serine residue of the N-box SLDD motif corresponds to the newly identified S8 phosphorylation site in *Sc*Yra1, raising the possibility that phosphorylation at this position directly modulates UBM-dependent protein interactions. The C-box of *Sc*Yra1 and *Ct*Yra1 additionally contains a putative SLED motif immediately upstream of the LDxxLD motif, which appears to be species-specific, as the corresponding sequence in ALYREF and *Sp*Mlo3 deviates more substantially from the UBM consensus.

To directly test whether the phosphorylation state of S8 influences Yra1 protein interactions, we purified N-terminally TAP-tagged Yra1 variants and analyzed co-purifying proteins by immunoblotting (Figures 4E and 4F and S4A–D). Despite the location of S8 within the N-box UBM, no significant change in Sub2 interaction is detected in any of the mutants tested. At 30°C, both TAP-yra1-S8A and TAP-yra1-S8A-S142A show an increased interaction with Hpr1 and Mex67 relative to wt Yra1 (Figure 4E). Under heat stress, the interaction of TAP-Yra1 with Hpr1 is strongly reduced regardless of the phosphorylation status (Figure S4C). Crucially, however, both TAP-yra1-S8A and TAP-yra1-S8A-S142A display a significantly increased interaction with Mex67 at 42°C compared to heat-stressed wt TAP-Yra1 (Figure 4F), demonstrating that phosphorylation of S8 alone is sufficient to weaken the Yra1-Mex67 interaction under heat stress. Consistent with S8 residing at a direct Mex67-binding interface, an AlphaFold multimer prediction^40^ of the Mex67-Mtr2 heterodimer in complex with Yra1 indicates that the N-terminal helix of Yra1, harboring the SLDD motif, directly contacts Mex67 and the C-terminus of Sub2 (Figures 4G and S4E–H).

Taken together, these results demonstrate that the SLDD motif in the N- and C-termini of *Sc*Yra1 each harbors a serine residue that undergoes stress-specific phosphorylation—at S8 during heat stress and at S211 during hyperosmotic stress. Preventing phosphorylation at S8 alone is sufficient to enhance Yra1-Mex67 interaction, identifying phosphorylation of Yra1-S8 as a mechanism that destabilizes the Yra1-Mex67 interface and thereby reduces Mex67 association with nuclear mRNPs and a concomitant inhibition of nuclear mRNA export during heat stress.

### Yra1-S8 phosphorylation is required for efficient binding to and expression of heat stress-induced transcripts

As described above, preventing phosphorylation at Yra1-S8 increases the interaction of Yra1 with Mex67, which in turn leads to enhanced association of the export receptor with nuclear mRNPs. To determine whether this altered protein interaction affects the RNA-binding properties of Yra1, we performed RNA immunoprecipitation (RIP) experiments with TAP-tagged Yra1 variants. Under heat stress, we compared binding of TAP-Yra1 and TAP-yra1-S8A to three highly expressed non-heat shock transcripts (*ILV5*, *CCW12*, *ACT1*) and two canonical heat shock transcripts (*SSA4*, *HSP104*) (Figures 5A and S5A). While TAP-yra1-S8A binding to non-heat shock transcripts remains unaltered, binding to heat shock transcripts is significantly reduced. No changes in RNA binding are detected in the *yra1-S8D* phosphomimetic mutant, which potentially mimics the heat-induced phosphorylation of Yra1 at 30°C (Figure S5B and C). Consistently, Mex67-TAP binding to heat shock transcripts is also reduced in *yra1-S8A* cells at 42°C (Figures 5B and S5D), indicating that phosphorylation of Yra1-S8 is required for efficient recruitment of both Yra1 and Mex67 to heat-induced transcripts.

**Figure 5.**
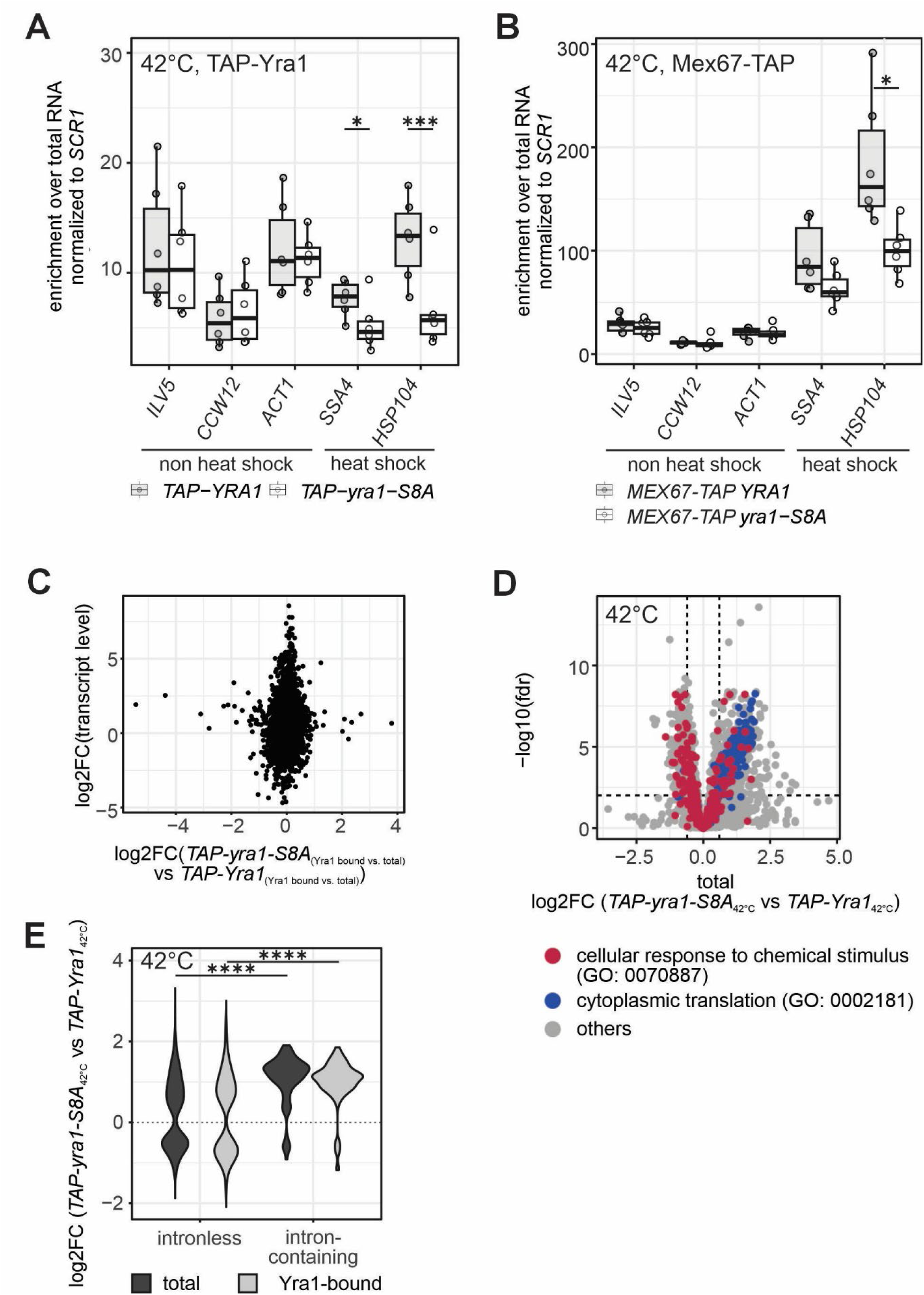
yra1-S8A binds less strongly to heat-induced transcripts and the expression of stress-induced mRNAs is decreased in *yra1-S8A* cells. (A and B) Binding of Yra1 (A) and Mex67 (B) to heat shock transcripts is reduced in *yra1-S8A* cells. RNA immunoprecipitations (RIP) from heat-shocked cells are quantified by RT-qPCR. CT values are normalized to the non-coding RNA *SCR1* and to the respective lysate sample (n ≥ 6). RIPs from cells grown at 30°C and representative Coomassie stained gels are shown in Figure S5. (C) The *yra1-S8A* mutation does not alter the intrinsic RNA-binding specificity of Yra1. When Yra1-bound RNA is normalized to total RNA, the correlation between transcript induction and differential binding of *TAP-yra1-S8A* is lost, indicating that the altered binding profile of the mutant is a consequence of changes in transcript levels rather than changes in RNA-binding specificity. Scatter plot of the log2 fold change of *TAP-yra1-S8A*-bound versus *TAP-YRA1*-bound RNA normalized to total RNA, plotted against the log2 fold change in transcript abundance upon a shift from 25°C to 37°C (RNA abundance t16 vs. t0 from).^41^ (D) The abundance of transcripts encoding proteins involved in the cellular response to chemical stimuli is reduced in *yra1-S8A* cells at 42°C. Volcano plot comparing total transcript levels in the *TAP-yra1-S8A* mutant versus *TAP-YRA1* cells. Transcripts annotated with the GO terms cellular response to chemical stimulus (GO 0070887, red) and cytoplasmic translation (GO 0002181, blue) are highlighted. (E) Intron-containing transcripts bind better to TAP-yra1-S8A (light grey) and are increased in total RNA (dark grey). Violin plot comparing the log_2_FC of TAP-yra1-S8A-normalized to TAP-YRA1-bound RNA for intron-containing and intronless transcripts. No difference between samples results in a log_2_FC of 0 (dotted line). ⃰⃰ ⃰⃰ ⃰⃰ p < 0.001, ⃰⃰ ⃰⃰ ⃰ ⃰⃰ p < 0.0001 by multiple unpaired t-test

To assess whether these findings reflect a global change in Yra1 RNA-binding specificity, we coupled the RIP experiment to RNA sequencing. Principal component analysis reveals good separation between total and Yra1-bound samples, with less pronounced but visible differentiation between wt and *yra1-S8A* mutant samples (Figure S5E). Plotting the binding ratio of TAP-yra1-S8A versus TAP-Yra1 against transcript induction upon heat shock^41^ reveals a slight but significant correlation: transcripts less efficiently bound by TAP-yra1-S8A tend to be more strongly induced by heat stress (Figure S5F). However, this correlation is lost upon normalization of Yra1-bound RNA to total RNA levels (Figure 5C), indicating that the reduced binding of yra1-S8A to heat shock transcripts reflects decreased expression of these transcripts in the mutant rather than an intrinsic change in RNA-binding specificity.

Consistent with this interpretation, comparison of total RNA from both strains reveals that transcripts associated with the response to chemical stress (GO:0070887) are significantly downregulated in *TAP-yra1-S8A* cells, while transcripts encoding proteins involved in translation (GO:0002181) are upregulated (Figures 5D and S5G). Furthermore, intron-containing transcripts are enriched in both the total RNA and the Yra1-bound fraction of *TAP-yra1-S8A* cells (Figure 5E), and an increase in intronic reads suggests an associated splicing defect in this mutant (Figure S6A).

Taken together, these results demonstrate that phosphorylation of Yra1 at S8 is required for efficient binding of both Yra1 and Mex67 to heat stress-induced transcripts and for the full induction of the heat stress transcriptional response. The reduced association of *yra1-S8A* with stress transcripts thus reflects impaired expression of heat-induced mRNAs rather than an altered intrinsic RNA-binding specificity of the mutant protein (Figure S6A).

### Yra1-S8 phosphorylation restricts nuclear export of heat shock transcripts

Our initial hypothesis was that yra1-S8A recruits more Mex67 to bulk mRNAs during heat shock, thereby driving increased export of transcripts that are normally retained in the nucleus. This would have explained the reduced nuclear poly(A)^+^ RNA accumulation observed by FISH. However, both the RIP RT-qPCR and sequencing data instead point to a specific role of Yra1-S8 phosphorylation in the synthesis and/or export of heat shock transcripts. To determine whether Yra1 and Mex67 exhibit similar binding preferences across the transcriptome, we plotted the enrichment of Yra1-bound transcripts normalized to total RNA against the corresponding Mex67-bound enrichment (Figure 6B). Transcripts more abundant after mild heat shock are highlighted in red.^41^ This analysis reveals that all mRNAs, including heat shock transcripts, display equivalent binding preferences for Mex67 and Yra1, indicating that neither factor selectively associates with heat-induced transcripts.

**Figure 6.**
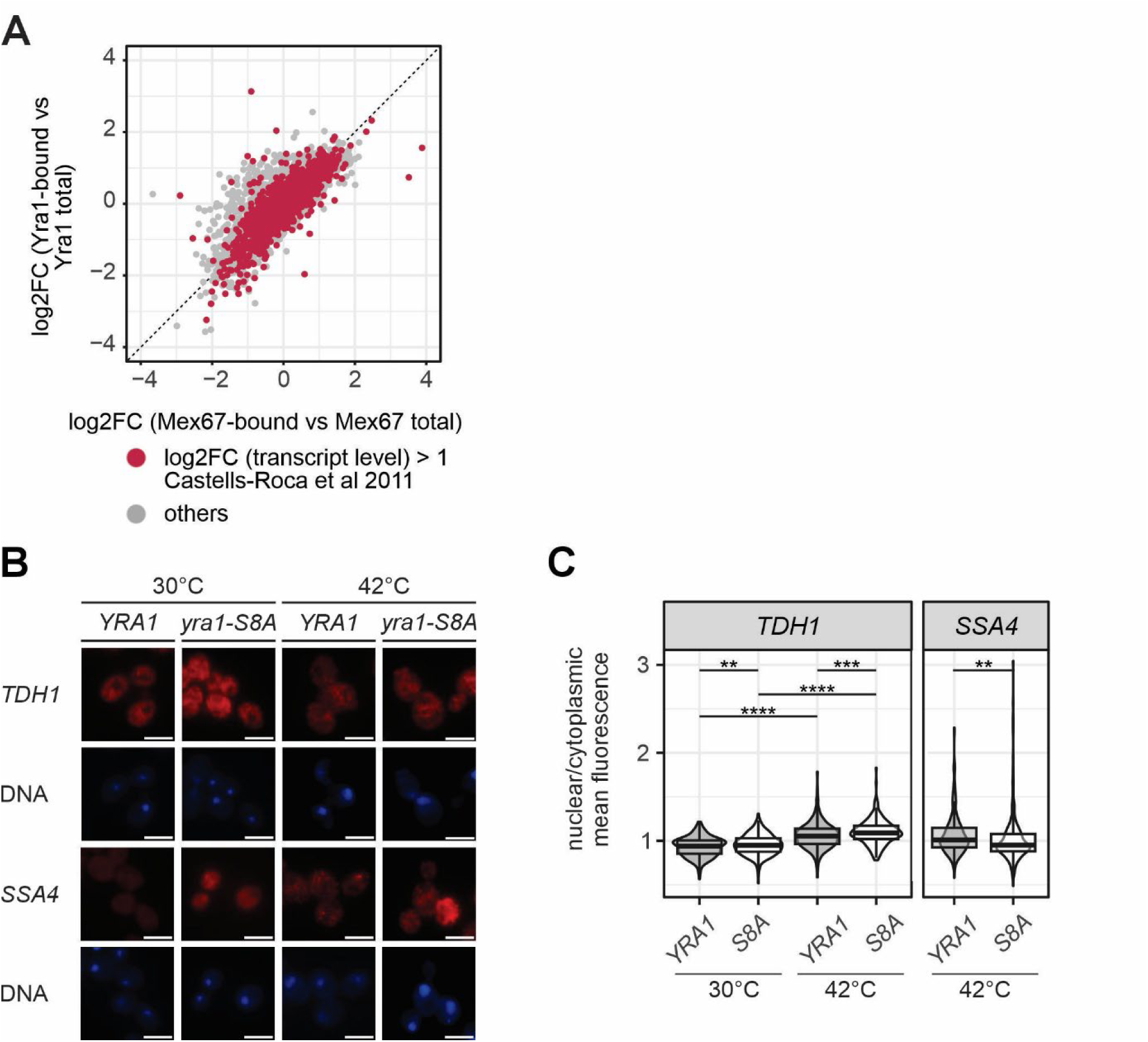
The *yra1-S8A* mutation leads to enhanced nuclear export of heat shock transcripts. (A) Binding to transcripts correlates between Yra1 and Mex67. Scatter plot showing the correlation between the log_2_FC enrichment (bound vs. total RNA) for transcripts bound to Yra1 and Mex67. Transcripts with a more than 2-fold increased abundance after shift from 25°C to 37°Care highlighted in red.^41^ (B) Nuclear export of the heat shock mRNA *SSA4* is enhanced in *yra1-S8A* cells at 42°C, while nuclear export of the *TDH1* mRNA is reduced. Representative smFISH images showing the localization of the heat shock transcript *SSA4* and the control transcript *TDH1* (encoding GAPDH) in *YRA1* and *yra1-S8A* cells at 30°C and 42°C. The Cy3-labeled probes detect the corresponding mRNA (red). DNA is stained with DAPI (blue). Scale bars: 5 µm. (C) Quantification of the nuclear-to-cytoplasmic signal ratios for the *SSA4* and *TDH1* mRNA as exemplified in panel (B). For each replicate at least 90 cells per sample were quantified (n = 3). Statistical significance: * p < 0.05, ⃰⃰ ⃰⃰ p < 0.01, ⃰⃰ ⃰⃰ p < 0.001, ⃰ ⃰⃰ p < 0.0001 by Welch t-test. For additional quantification of the total Cy3 signals see Figure S6.

Since static RIP binding data cannot fully capture the dynamics of nuclear mRNA export, we use smFISH to directly assess the effect of the *yra1-S8A* mutation on the nuclear export of the heat shock transcript *SSA4* (Figures 6B and 6C). Strikingly, after 1 hour of heat stress, the nuclear-to-cytoplasmic fluorescence ratio of *SSA4* is significantly reduced in *yra1-S8A* cells compared to wt (Figures 6B and 6C), indicating that heat shock transcripts are more efficiently exported when S8 phosphorylation is prevented. To assess the specificity of this effect, we additionally examine the distribution of *TDH1* (encoding GAPDH), a non-heat shock transcript. Although the overall *TDH1* fluorescence signal decreases after 1 hour of heat stress, the nuclear-to-cytoplasmic ratio of *TDH1* increases in both strains, consistent with its known nuclear retention during heat shock (Figures 6C and S6C). Notably, this nuclear retention of *TDH1* is further enhanced in *yra1-S8A* cells relative to wt controls (Figure 6C). This demonstrates that the *yra1-S8A* mutation does not globally promote mRNA export but rather has opposing effects on distinct transcript classes: it enhances export of heat shock transcripts while further restricting export of non-heat shock transcripts.

Taken together, these findings indicate that phosphorylation of Yra1 at S8 restricts the nuclear export of heat shock transcripts. The reduced nuclear poly(A)^+^ RNA accumulation observed in *yra1-S8A* cells therefore reflects enhanced export of heat shock transcripts rather than increased export of bulk mRNA. Since *SSA4* transcript levels are not altered in *TAP-yra1-S8A* cells (Figures S6A and S6C), reduced transcription is unlikely to account for this observation. We therefore propose the following model: under heat stress, Slt2-dependent phosphorylation of Yra1 at S8 promotes the synthesis of heat-induced transcripts while simulataneously weakening the Yra1-Mex67 interaction, thereby reducing Mex67 recruitment to nuclear mRNPs and limiting nuclear export of these trancripts.

## DISCUSSION

### Nuclear mRNA accumulation is a stress-specific rather than a universal response

Adaptation to environmental change including during development or disease states is essential for cell survival. While short-term responses rely heavily on post-translational modifications, lasting adaptation requires changes in gene expression that enable long-term stress survival.^42^ Stress responses range from generic to highly stress-specific^43^, and our data demonstrate that nuclear mRNA accumulation belongs to the latter category: it is not triggered by all stress conditions and therefore does not represent a universal stress response. Strikingly, certain conditions—rapamycin, arsenite and DTT treatment—activate Slt2 and promote Nab2 phosphorylation without causing any detectable nuclear mRNA retention, demonstrating that Slt2 activation and Nab2 phosphorylation are not sufficient to drive nuclear mRNA accumulation. Conversely, nutrient starvation triggers nuclear mRNA retention independently of both Slt2 and Nab2 phosphorylation, while hyperosmotic stress causes nuclear mRNA accumulation accompanied by Nab2 phosphorylation through an Slt2-independent mechanism. These observations reveal an unexpected complexity in the regulation of nuclear mRNA export and suggest that multiple mechanistically distinct pathways converge on nuclear mRNA retention in a stress-specific manner. Dissecting these pathways will be an important goal for future studies.

The identity of the transcripts that accumulate in the nucleus during stress also remains incompletely understood. An increase in nuclear *GAPDH/TDH1* mRNA upon heat stress is consistent with the retention of bulk mRNA, as proposed previously.^1^ However, our data challenge this view: rather than causing increased export of bulk mRNA, preventing Yra1-S8 phosphorylation specifically enhances nuclear export of heat shock transcripts while further restricting export of the non-heat shock transcript *TDH1*, suggesting that heat shock transcripts themselves contribute to the nuclear poly(A)^+^ RNA pool detected by oligo(dT) FISH. These observations highlight a current limitation of the field: the precise composition and functional significance of nuclear-retained mRNAs during stress remain poorly defined.^39^ Additional contributing factors, such as stress-induced changes in polyadenylation or mRNA turnover, cannot be excluded. The hypothesis that bulk mRNA is selectively retained to prioritize ribosome engagement with heat shock transcripts is conceptually attractive; however, *Δslt2* cells, which lack heat shock-induced nuclear poly(A)^+^ RNA accumulation, still produce wild-type levels of heat shock proteins^37^, arguing against this model in its simplest form. The exact composition of stress-retained mRNAs, their function during stress adaptation and the full mechanistic basis of their nuclear retention warrant further investigation.

### Stress-specific remodeling of the mRNA export factor phosphoproteome

Our phosphoproteomic analyses reveal widespread and highly stress-specific remodeling of the phosphorylation landscape of nuclear mRNA export factors, with heat stress and hyperosmotic stress producing clearly distinct phosphorylation signatures on proteins annotated to the mRNA export pathway. This stress specificity is consistent with the engagement of mechanistically distinct signaling pathways and provides a molecular basis for the observed differences in mRNA export regulation between these two conditions. Post-translational modifications are well-established regulators of nuclear mRNA export: phosphorylation of Npl3 at S411 drives its nuclear import, while subsequent dephosphorylation by Glc7 enables RNA binding and Mex67 recruitment.^25,44^ Ubiquitination of Hpr1 stabilizes its interaction with Mex67^26,45^, while Tom1-dependent ubiquitination of Yra1 triggers its dissociation from nuclear mRNPs prior to nuclear pore complex (NPC) transit.^24^ Our data add phosphorylation to the repertoire of modifications that regulate Yra1 function and suggest an interplay between phosphorylation and ubiquitination at the Yra1 N-terminus: the Rsp5-dependent ubiquitination site K7 lies immediately adjacent to the S8 phosphorylation site identified here^24,46^, raising the intriguing possibility that these two modifications are mutually regulated or functionally coupled. Notably, the reduced protein levels observed in the *yra1-S8A-S142A* mutant hint at a change in ubiquitination, although no direct correlation between Yra1 phosphorylation and ubiquitination could be established (data not shown). Since the absence of Yra1 phosphorylation affects the association of Mex67 and Sub2 with nuclear mRNPs rather than that of Yra1 itself, phosphorylation and ubiquitination likely act at distinct steps of the mRNA export cycle.

Strikingly, despite the large number of differentially phosphorylated residues identified across the mRNA export machinery, mutation of a single amino acid—S8 of Yra1—is sufficient to significantly alter nuclear mRNA export during heat stress. This highlights the disproportionate regulatory importance of individual phosphorylation events within otherwise complex signaling networks and underscores the value of unbiased phosphoproteomic approaches for identifying functionally critical modifications. However, the significant but partial reduction in nuclear mRNA accumulation caused by *yra1-S8A* mutation suggests that additional Slt2 targets and/or parallel regulatory mechanisms contribute to the full extent of heat stress-induced mRNA export inhibition. Interestingly, the C-terminus of Hpr1 is not only phosphorylated during heat stress but also sumoylated during acid stress, a modification required for the binding and stabilization of acid stress-induced transcripts^47^, suggesting that Hpr1, like Yra1, may play a role in the selective handling of stress-induced transcripts. The combination of *yra1-S8A* with *nab2* phosphorylation site mutants does not further reduce nuclear mRNA accumulation relative to the single mutants (data not shown), suggesting partial epistasis within the Slt2-dependent pathway. Fully resolving this regulatory network will require systematic epistatic analyses and the identification of additional Slt2 substrates within the mRNA export machinery. The phosphoproteomic dataset generated here provides a valuable resource for such future investigations. Combined with the phosphoproteomic data comparing a variety of stress conditions will allow a comprehensive analysis of the regulatory networks.^48^

### Phosphorylation of the Yra1 N-terminal UBM as a checkpoint for Mex67 recruitment and nuclear mRNA export

The identification of Yra1-S8 phosphorylation within its conserved N-terminal SLDD/E UBM provides important mechanistic insight into how Yra1 phosphorylation regulates nuclear mRNA export. The UBMs of Yra1/ALYREF have recently come into focus as key mediators of mRNP assembly and remodeling: in human cells, the WxHD motif of ALYREF mediates mRNP compaction through interaction with the EJC, while the LDxxLD motif stimulates UAP56 ATPase activity and drives mRNP remodeling.^18–21^ Yra1 is the most prominent protein component of highly compacted yeast nuclear mRNPs^23^, and the “IDR network model” posits that Yra1, through its intrinsically disordered regions, drives RNA-RNA interactions and mRNP compaction.^23^ Our AlphaFold-based structural prediction indicates that the N-terminal helix of Yra1, harboring the SLDD motif, directly contacts Mex67, providing a structural rationale for how S8 phosphorylation destabilizes the Yra1-Mex67 interface. We therefore propose that under normal growth conditions, the unphosphorylated SLDD motif of Yra1 promotes Mex67 recruitment to the nuclear mRNP, an interaction required for mRNP export competence.^11,49^ During heat stress, Slt2-dependent phosphorylation of S8 weakens this interaction, reducing Mex67 association with nuclear mRNPs and thereby limiting their nuclear export. Yra1 dissociation from the mRNP may subsequently be triggered by Tom1-dependent ubiquitination, as described previously.^24^ This model is consistent with the temperature-sensitive *GFP-yra1-8* mutant, which carries D10E11-to-K10K11 substitutions within the same N-terminal SLDD/E motif and displays nuclear retention of several transcripts including heat shock mRNAs, as well as enhanced interaction with Mlp2 and strongly reduced Mex67-Mlp2 interaction^50,51^ —a phenotypic profile strikingly similar to that observed here for *yra1-S8A*.

Whether the regulatory function of UBM phosphorylation is conserved across species is an important open question. Our data reveal stress-specific phosphorylation at S8 (heat stress) and S211 (hyperosmotic stress) within the N- and C-terminal SLDD/E motifs of *S. cerevisiae* Yra1, respectively, and phosphorylation at equivalent positions has been detected in phosphoproteomic datasets from other organisms.^52^ In mammalian cells, Slt2 is functionally homologous to ERK1/2 and ERK5, both of which are activated by environmental stress and regulate gene expression at multiple levels.^53^ ERK-dependent phosphorylation of human ALYREF has not been described, but given the conservation of the SLDD/E motif and its regulatory importance demonstrated here, it is tempting to speculate that analogous phosphorylation events modulate ALYREF-NXF1 (the human Mex67 orthologue) interaction and mRNA export efficiency during stress in higher eukaryotes. Whether this mechanism extends to other UBM-containing proteins such as UIF and LUZP4^21^ represents another avenue for future investigation.

### Yra1-S8 phosphorylation coordinates heat shock transcript expression and selective nuclear mRNA export

Perhaps the most unexpected finding is that preventing Yra1-S8 phosphorylation not only alters mRNA export but also impairs the expression of heat stress-induced transcripts and efficient splicing. The RNA-binding specificity of *yra1-S8A* is not intrinsically altered—normalization of Yra1-bound RNA to total RNA eliminates the apparent difference in heat shock transcript binding—indicating that the reduced association of yra1-S8A with stress-induced mRNAs is a secondary consequence of their reduced expression rather than a direct effect on RNA-binding activity. This points to an unexpected role of Yra1-S8 phosphorylation in the transcriptional or co-transcriptional phase of heat shock gene expression. While the THO complex, which physically interacts with Yra1, is required for transcription elongation, THO recruitment itself is not dependent on Yra1.^10^ Nevertheless, Yra1 is recruited to intron-containing genes, and Yra1 itself harbors a functionally important intron that regulates Yra1 protein levels.^54^ The increased intronic read abundance detected in *TAP-yra1-S8A* cells suggests a splicing defect that may reflect impaired Sub2 function or defective mRNA surveillance, given the known connections between Yra1/ALYREF, Sub2/UAP56 and the splicing machinery.^11,21^ In addition, ALYREF and TREX are involved in sensing transcription inhibition and storage of export competent and NXF1-bound transcripts in nuclear speckles in human cancer cells, further strengthening the link between ALYREF/Yra1 and transcription.^55^ Further studies will be required to determine whether Yra1-S8 phosphorylation directly influences transcription, splicing or mRNA surveillance.

Taken together, our findings support a model in which Slt2-dependent phosphorylation of Yra1-S8 during heat stress serves a dual function: it promotes the expression of heat-induced transcripts while simultaneously weakening the Yra1-Mex67 interaction, reducing Mex67 recruitment to nuclear mRNPs and thereby limiting the nuclear export of heat shock mRNAs. The net consequence is a paradoxical situation in which the cell actively restricts export of the very transcripts it has just induced. One possible interpretation is that transient nuclear retention of heat shock mRNAs serves a quality control function, ensuring that only fully processed and properly assembled mRNPs are exported and translated during the acute stress phase. Alternatively, nuclear retention may provide a means of rapidly adjusting the cytoplasmic availability of heat shock mRNAs in a switch-like manner upon stress relief, when Slt2 activity and Yra1 phosphorylation are reversed. Resolving the physiological significance of this regulatory mechanism and determining whether it is conserved in the mammalian ERK-ALYREF-NXF1 pathway, will be important directions for future work.

## RESOURCE AVAILABILITY

### Lead contact

Requests for further information and resources should be directed to and will be fulfilled by the lead contact, Katja Sträßer.

### Materials availability

Strains and plasmids generated in this study will be made available on request.

## ACKNOWLEDGMENTS

We thank Vera Bettenworth, Daniel Bauer and Nils Holger Maier for critical reading of the manuscript. We are grateful to Cornelia Kilchert for her advice on the FISH quantification script. We thank our students Elena Bagrin and Cora-Lee Weber for preliminary experiments. This work was supported by the European Union (European Research Council Consolidator Grant, grant number 772049 to K.S.) and the German Research Foundation (RTG2355 and under Germanýs Excellence Strategy – EXC 2026, Cardio-Pulmonary Institute, Project ID: 390649896 to K.S.).

## AUTHOR CONTRIBUTIONS

Conceptualization, J.F.S. and K.S.; methodology, J.F.S.; MS data acquisition and analysis, J.G.; Investigation, J.F.S., J.D. and J.G.; writing—original draft, J.F.S..; writing—review & editing, J.F.S and K.S..; funding acquisition, K.S.; resources, K.S.; supervision, K.S..

## DECLARATION OF GENERATIVE AI AND AI-ASSISTED TECHNOLOGIES

During the preparation of this work, the authors used kiChat (Claude Sonnet 4.6) provided by the JLU for language editing. After using this tool or service, the authors reviewed and edited the content as needed and take full responsibility for the content of the publication.

## SUPPLEMENTAL INFORMATION

**Document S1. Figures S1–S6 and Table S1-S4**

## METHODS

### Yeast strains, plasmids and primers

Lists of all yeast strains, plasmids and primers used in this study can be found in the supplement. All plasmids created in this study were cloned using Gibson Assembly. For the used phosphorylation site mutants point mutations were introduced into a pRS315-*YRA1* plasmid. Afterwards, the plasmids were transformed into a *YRA1* shuffle strain and the pRS316-*YRA1* plasmid was shuffled out by restreaking twice on 5’FOA plates.

### Growth conditions

If not indicated otherwise yeast cells were inoculated to an OD_600_ of 0.2 and grown to an OD_600_ of 0.8 in YPD (10 % yeast extract, 20 % peptone, 20 % glucose) at 30°C, 180 rpm. The different stress conditions tested are listed in Table S1.

### Oligo(dT) fluorescent in situ hybridization

FISH to visualize the distribution of poly(A)^+^ RNA in the cell was performed as described in Keil et al. ^(^2023)^.58^ Briefly, cells were grown to an OD_600_ of 0.8 and if indicated stress condition was performed. Cells were fixed using 4 % formaldehyde, washed and spheroblased by digesting the cell wall using 100T Zymolyase. After washing, the spheroplasts were attached to ploy-lysine coated cover slips, prehybridized at 37°C for 1 h in prehybridization buffer (50% formamide, 10% dextran sulphate, 125 μg/ml of *E. coli* tRNA, 500 μg/ml hering sperm DNA, 4x SSC, 0.02% polyvinyl pyrrolidone, 0.02% BSA, 0.02% Ficoll-40) and incubated with 0.75 µl of 1 pmol/µl Oligo(dT)-Cy3 (50xT) overnight. After washing, the coverslips were mounted with ROTI Mount FluorCare DAPI (Carl Roth) on slides. Microscopy was performed using a DeltaVision Ultra High-Resolution microscope (Cytiva) with 60x magnification. Z-stack images were taken with a 0.2 µm spacing. Image quantification was performed in FIJI ImageJ using a semiautomated script.^57^ The cells were identified by thresholding of the Oligo(dT)-Cy3 signal and the nuclei using the DAPI signal. The average intensity of the Oligo(dT)-Cy3 signal was measured in the whole cell, cytoplasm and nucleus using average intensity Z-projections. The ratio of mean nuclear-to-cytoplasmic fluorescence was calculated and visualized in R.^59^

### Phosphoproteome

For each sample, 400 ml yeast culture were grown to mid log phase and stress treated as indicated. The cells were resuspended in 1 ml RNA IP buffer (25 mM TrisHCl pH 7.5, 150 mM NaCl, 2 mM MgCl2, 0.2% Triton X-100, 500 µM DTT) containing protease inhibitor and phosphatase inhibitor (PhosStop, Roche) and lysed using a FastPrep-24 5G machine (3x 20 sec at 6 m/s). The lysate was cleared by sequential centrifugation steps at 4000 rpm and 13000 rpm, respectively, for 5 min at 4°C. Scientific Service Group Biomolecular Mass Spectrometry at the Max Planck Institute of Heart and Lung Development (Bad Nauheim, Germany). For each condition and replicate lysate equating to 1 mg total protein content was processed as described^60^, using MARMoSET^61^ for documentation of instrumentation, as well as peptide/spectrum matching (see supplementary material) and autonomics (DOI: 10.18129/B9.bioc.autonomics) for downstream bioinformatic analysis.

### Quantitative Western blots

Yeast cells were grown to an OD_600_ of 0.8 and if indicated stressed. The cells were resuspended in 500 µl water, mixed with 150 µl pretreatment solution (1.85 M NaOH, 7.5 % beta-mercaptoethanol) and incubated for 20 min on ice. Afterwards, 82.5 µl 100 % TCA was added, incubated and centrifuged for precipitation. The supernatant was removed and the pellet resuspended in SDS-loading dye with additional 1 M Tris. The samples were boiled for 10 min, separated on an SDS-gel and transferred to nitrocellulose membrane. After blocking of the membrane, the proteins were detected using the appropriate primary antibodies and incubated with a secondary horse radish peroxidase coupled antibody. The signal was detected using Clarity Western ECL Substrate solution (Biorad) and the ECL ChemoCam Imager (Intas). Signals were quantified using FIJI ImageJ.^57^ The antibody used for Cbp80 detection is a kind gift of D. Görlich.^62^Click or tap here to enter text.

### TAP purification

Protein purifications were performed with a TAP tag consisting of a calmodulin-binding peptide (CBP), a TEV cleavage site and protein A. To determine the composition of nuclear mRNPs a one-step TAP purification using Cbp20-TAP was performed. Interaction partners of TAP-Yra1 were purified using a two-step TAP purification.

For Tandem Affinity Purification^63^ 2 L TAP-tagged yeast cells were grown to an OD_600_ of 3.3-3.9. After harvesting, the pellet was resuspended in 1.5 ml ice-cold 1xPBS and flash frozen in liquid nitrogen. The cell suspension beads were lyses mechanically in liquid nitrogen using the freezer mill 6870D (SPEX SamplePrep). Cell powder was resuspended in TAP buffer (100 mM NaCl, 50 mM HEPES pH 7.5, 1.5 mM MgCl_2_, 0.15 % NP-40, 1 mM DTT) containing either protease inhibitor (30°C samples) or protease inhibitor and phosphatase inhibitor (42°C samples). The lysate was cleared by subsequent centrifugation at 3488 g and 16500 g for 12 and 60 min, respectively. The supernatant was incubated with 600 µl IgG beads slurry (IgG Sepharose 6 Fast Flow, Cytiva) for 1.5 h at 4°C. The beads were washed and incubated with TEV protease to elute the Calmodulin-binding peptide tagged protein from the beads. If a two-step purification was performed, the eluate was mixed with Calmodulin Affinity Resin (Agilent) preincubated with 4 mM CaCl_2_ and incubated for 1 h at 4°C. The beads were washed and the bound proteins eluted using EGTA elution buffer (10 mM HEPES pH 8, 25 mM EGTA pH 8) for 15 min, 1000 rpm at 37°C. The eluted proteins were TCA precipitated before Western blot analysis. Copurified proteins were analyzed using Western blot and quantified using FIJI ImageJ.^57^

### RNA-immunoprecipitation (RIP)

RIP was performed similar as described in Keil et al. (2023).^58^ Briefly, the TAP-tagged yeast strains were grown in YPD to an OD_600_ of 0.8. Cells were harvested, flash frozen and stored at −80°C. To perform RIP experiments the samples were thawed on ice and resuspended in RIP buffer (25 mM Tris–HCl pH 7.5, 150 mM NaCl, 2 mM MgCl2, 0.2% Triton X-100, 500 µM DTT) and protease inhibitor. The cells were lysed using the FastPrep-24 5G and glass beads. The lysate was cleared by centrifugation and DNA was digested using DNaseI (Thermo Fisher Scientific) for 30 min on ice. For each sample 30 µl pre-washed IgG-coupled Dynabeads (Thermo Fisher Scientific) were added and incubated for 3 h at 4°C. The beads were washed with RIP buffer and RNA was isolated using TRIZOL (Thermo Fisher Scientific). After removing the aqueous phase containing the RNA, the remaining sample was acetone precipitated to analyze protein purification. The RNA in lysate and eluate was analyzed by reverse transcription followed by RT-qPCR. For each RNA the enrichment was calculated over the lysate and the signal of the small, cytoplasmic ncRNA *SCR1*.

For heat shock samples only 300 ml cells were cultured and mixed with 100 ml hot media to induce the heat shock. Heat shock was performed for 15 min at 42°C to ensure the high expression of heat shock mRNAs. The RIP buffer used for heat shock RIPs phosphatase inhibitor was added and the lysate was incubated for 1.5 h at 4°C instead of 3 h.

### RIP sequencing and analysis

The RIP was performed as described above. The RNA of lysate and eluate was isolated using hot phenol RNA isolation. Residual DNA was removed by DNaseI (Thermo Fisher Scientific) digest for 1 h at 37°C. Afterwards, RNA was isolated using phenol. Library preparation, sequencing and initial data processing including alignment to the sacCer3 reference genome was performed at the GeneCore EMBL. The samples were processed with a Biomek i7 Handaling system following the NEB UltraII Directional mRNA workflow after manufacturer’s instructions with a fragmentation time of 6 min. An adapter dilution of 1 to 30 was used and the library amplified using NEB UDIs. Sequencing was performed using the Illumina Nextseq 2000 using a P2-100cycle kit. Alignment was performed using STAR.^64^ To determine the fold changes between the samples the DESeq2 tool after read counting with HTSeq-count in Union mode on the GCF_000146045.2_R64 (scaCer3) gene model was used on the Galaxy server.^65–67^ Additional Deseq2 analysis was performed in R^59^ after HTSeq count on the Galaxy server using the GSE188455_Saccharomyces_cerevisiae.EF4.68_SGDv64 gene model to determine binding to antisense RNAs.^68^ All further analysis was performed with R and visualized with the ggplot2 package. GO term annotations were loaded using biomaRt.^69^ Furthermore the tidyverse, dplyr, ggpubr and ggrepel packages were used.^66,70–74^

### Single molecule FISH

Cells were inoculated to an OD_600_ of 0.1 and grown to 0.4. Heat shock was performed for 60 min. Cells were subsequently crosslinked with 3.6 % formaldehyde for 25 min and 20 min at 42°C and 30°c, respectively. Cells were harvested, washed with smFISH buffer (1.2 M sorbitol, 0.1 M potassium phosphate dibasic pH 7.5) and treated with 100T Zymolyase. After spheroblasting, cells were attached to poly-lysine coated coverslips and incubated in 70 % ethanol overnight at 4°C. Slides were removed from ethanol, dried and incubated at 30°C overnight with 50 µl hybridization buffer (10 % dextran sulphate, 0.1 % *E. coli* tRNA, 2x SSC, 0.02 % BSA, 10 % Formamide, 2 mM Vanadyl Ribonucleoside Complex) containing 125 nM *SSA4* smFISH probe (Stellaris, LGC Biosearch Technologies). The slides were washed with buffer A (2x SSC, 10 % Formamide) and 1x PBS for 30 min at 30°C and 2 min at room temperature, respectively. The dry coverslips were mounted and microscopy was performed as described previously for the Oligo(dT) FISH.

Image quantification was performed with the same FIJI ImageJ script used for analysis of the Oligo(dT) FISH.^57^ Since the *SSA4* mRNA is not as evenly distributed across the whole cell as all poly(A)^+^ RNA in the Oligo(dT) FISH defining the cell border with Cy3 background signal was more difficult. Therefore, the intensity maxima in the nucleus were counted, measured and compared between the samples to prevent excluding cells with very strong nuclear accumulation in the analysis. The measurements were visualized in R.

### Quantification and statistical analysis

Statistical tests were performed to determine significance between samples using either R or GraphPad Prism. The number of replicates can be found in the corresponding figure legends.

