## Supplemental Data for "Phosphorylation of the mRNP component Yra1 couples heat stress to nuclear mRNA export inhibition"

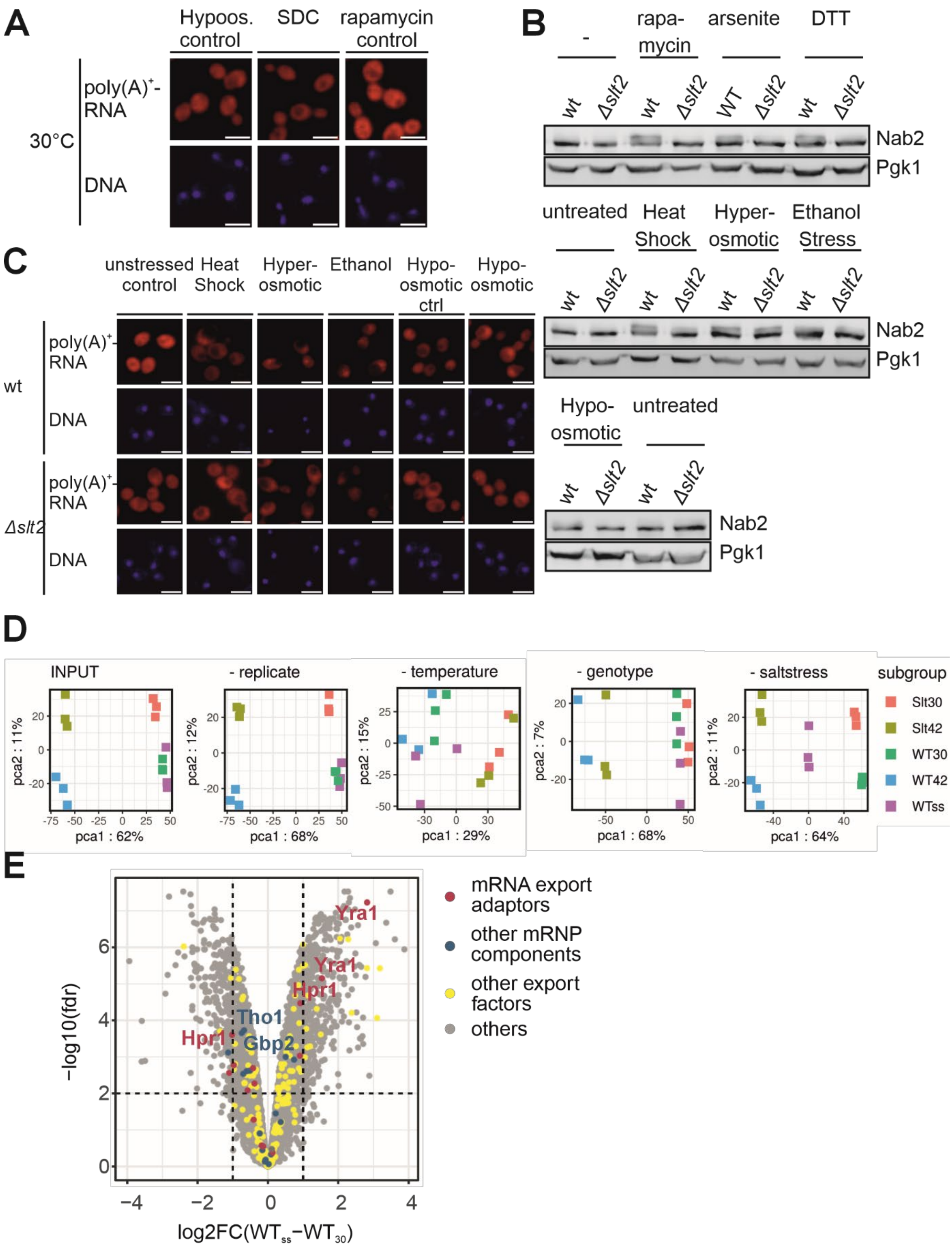

**Figure S1. Nuclear poly(A)$^+$ RNA accumulation and phosphorylation of mRNA export proteins are stress-specific, related to Figure 1**

(A) Nuclear accumulation of poly(A)$^+$ RNA occurs under heat, hyperosmotic, ethanol and hypoosmotic stress as well as glucose and nitrogen starvation, but not in response to other stress conditions. Representative microscopy images of cells exposed to different stress conditions, analysed by oligo(dT)-Cy3 FISH to determine the localization of poly(A)$^+$ RNA (red). Nuclear DNA is stained with DAPI (blue). Scale bars: 5 µm.

(B) Phosphorylation of the mRNP component Nab2 and its kinase Slt2 is stress specific. Nab2 and Slt2 phosphorylation under the indicated stress conditions is assessed in wt and *Δslt2* cells by Western blotting. Pgk1 serves as a loading control.

(C) Slt2-depenent nuclear mRNA accumulation occurs under heat, ethanol and hypoosmotic stress. Representative microscopy images of cells exposed to different stress conditions, analysed by oligo(dT)-Cy3 FISH to determine the localization of poly(A)$^+$ RNA (red). Nuclear DNA is stained with DAPI (blue). Scale bars: 5 µm.

(D) Replicates of the phosphoproteome samples cluster together in Principle component analysis (PCA).

(E) Phosphorylation of nuclear mRNA export factors is dynamically regulated in response to different stresses. Volcano plot depicting the $log_2$-fold changes (log2FC) in the phosphorylation level of different amino acids in wt cells exposed to hyperosmotic (salt) stress (ss) compared to untreated wt cells (30°C), plotted against the negative $log_{10}$ of the false discovery rate (FDR). Proteins involved in nuclear mRNA export are colored in red, petrol and yellow for export adaptors (Nab2, Npl3, Yra1, Hpr1), additional mRNP components (other TREX components, Tho1, CBC) and additional proteins associated with the GO term “mRNA export from nucleus” (GO:0006406 (2021), respectively.

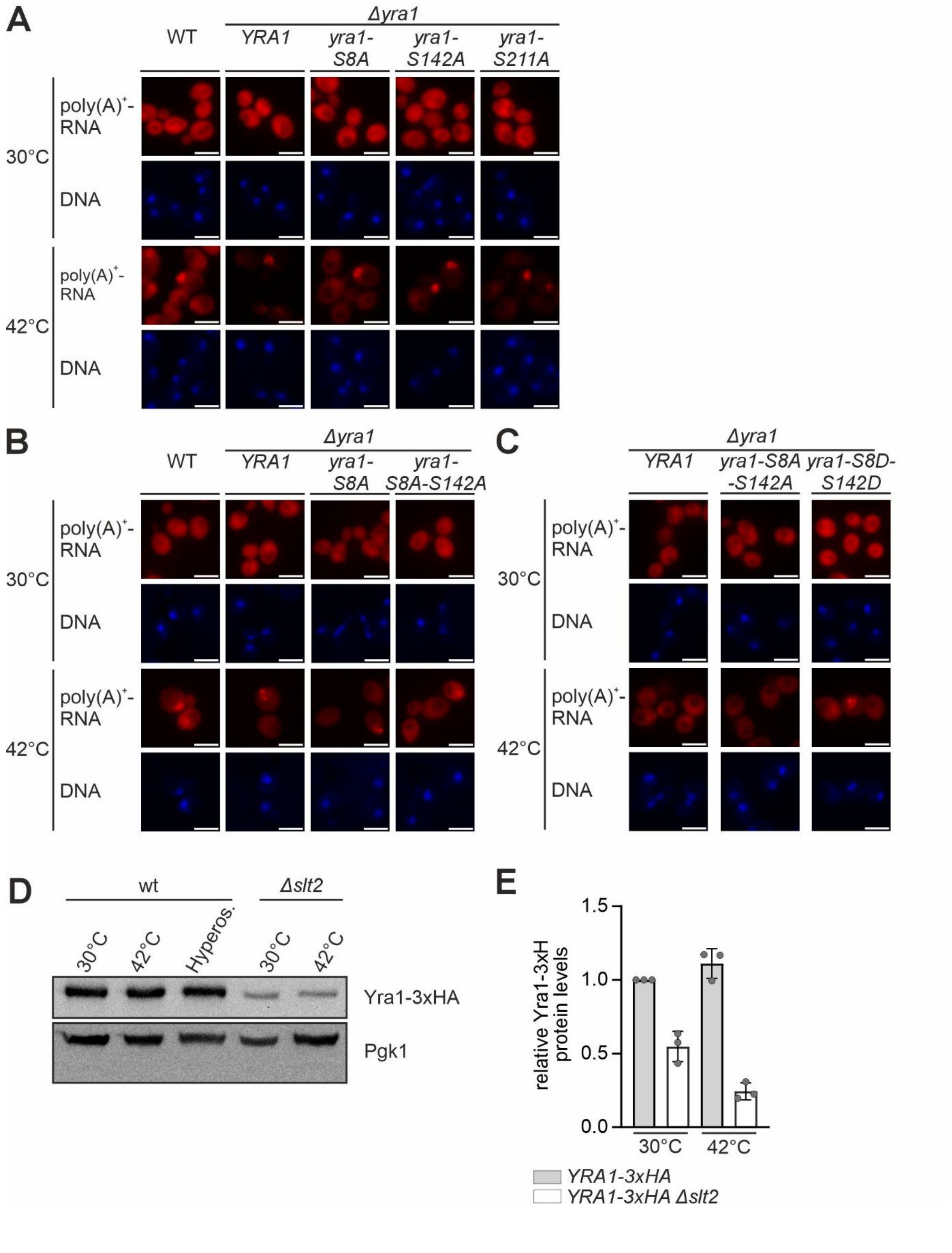

A
WT
Δyra1
YRA1
yra1-S8A
yra1-S142A
yra1-S211A
poly(A)+-RNA
DNA
30°C
42°C
B
Δyra1
yra1-S8A-S142A
C
yra1-S8A-S142A
yra1-S8D-S142D
D
wt
Δslt2
30°C
42°C
Hyperos.
Yra1-3xHA
Pgk1
E
relative Yra1-3xH protein levels
1.5
1.0
0.5
0.0
YRA1-3xHA
YRA1-3xHA Δslt2

**Figure S2. Phosphorylation of the mRNA export adaptor Yra1 at serine 8 (S8) in is critical for efficient heat stress-induced nuclear, related to Figure 3 and 4**

(A to C) Mutation of serine 8 to alanine (*yra1-S8A*), which prevents phosphorylation at this site, reduces the nuclear accumulation of poly(A)$^+$ RNA in response to heat shock. Representative microscopy images of cells exposed to different stress conditions, analysed by oligo(dT)-Cy3 FISH to detect poly(A)$^+$ RNA (red). Nuclear DNA is stained with DAPI (blue). Scale bars: 5 µm.

(D) Yra1-3xHA protein levels are reduced in *Δslt2* cells. Representative Western bots of total Yra1-3xHA protein levels in whole cell lysates of wt and *Δslt2* cells. Pgk1 serves as loading control. Consistent with this observation, a reduction in Yra1 abundance upon deletion of *SLT2* is also detected in the phosphoproteomic analyses.

(E) Quantification of the experiment shown in (D).

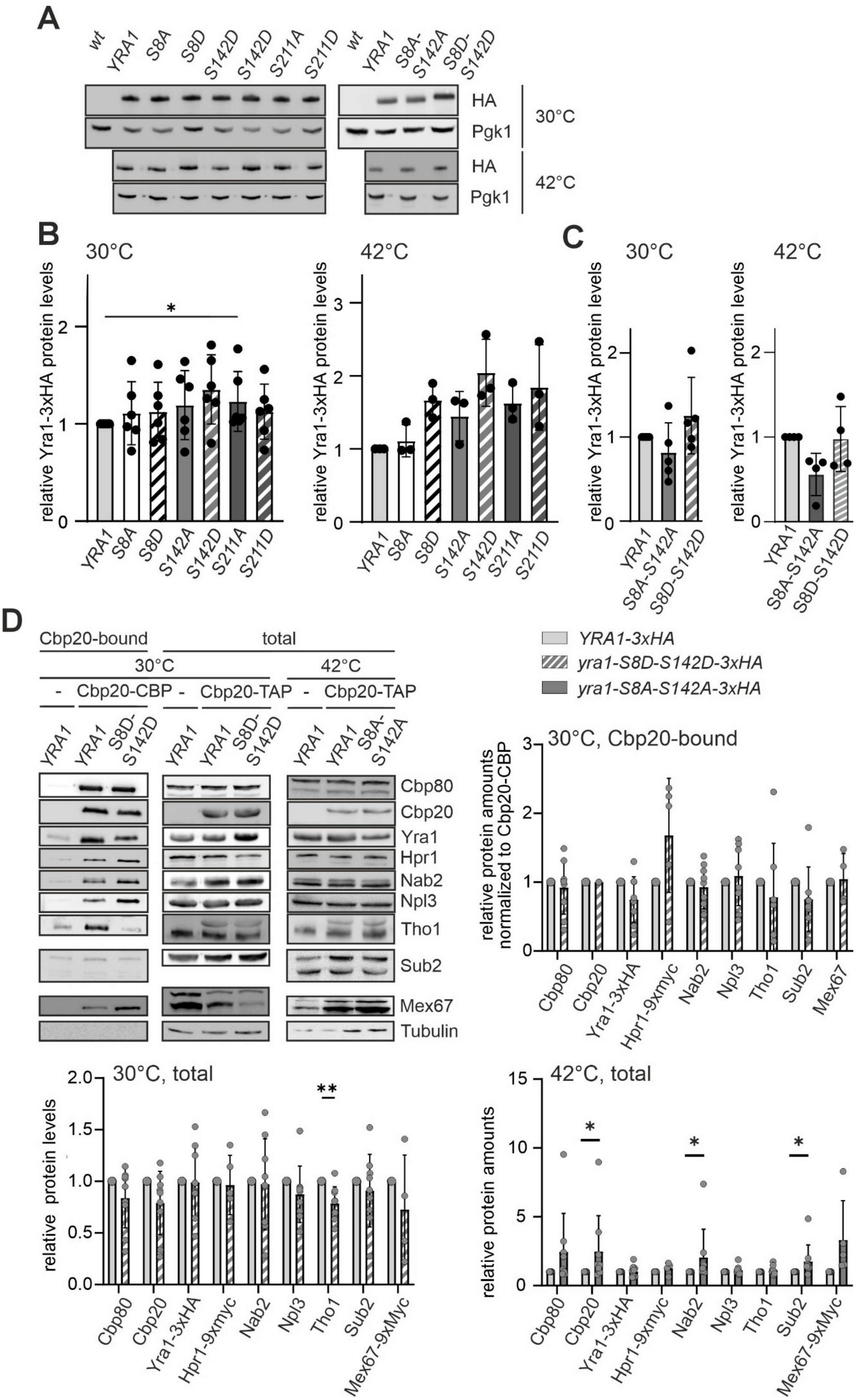
A
wt
YRA1
S8A
S8D
S142D
S142D
S211A
S211D
wt
YRA1
S8A-S142A
S8D-S142D
HA
Pgk1
30°C
HA
Pgk1
42°C
B
30°C
42°C
relative Yra1-3xHA protein levels
YRA1
S8A
S8D
S142A
S142D
S211A
S211D
C
30°C
42°C
relative Yra1-3xHA protein levels
YRA1
S8A-S142A
S8D-S142D
D
Cbp20-bound
total
30°C
42°C
Cbp20-CBP
Cbp20-TAP
Cbp20-TAP
YRA1
S8D-S142D
S8A-S142A
Cbp80
Cbp20
Yra1
Hpr1
Nab2
Npl3
Tho1
Sub2
Mex67
Tubulin
YRA1-3xHA
yra1-S8D-S142D-3xHA
yra1-S8A-S142A-3xHA
30°C, Cbp20-bound
relative protein amounts normalized to Cbp20-CBP
Cbp80
Cbp20
Yra1-3xHA
Hpr1-9xmyc
Nab2
Npl3
Tho1
Sub2
Mex67
30°C, total
relative protein levels
Mex67-9xMyc
42°C, total
relative protein amounts

**Figure S3. A reduction in Yra1 protein levels is observed exclusively in the *yra1-S8A-S142A* double mutant at 42°C, related to Figure 4**

(A to C) The *yra1-S8A-S142A* double but not the *yra1-S8A* single mutant has reduced Yra1 protein levels. (A) Representative Western blots of total Yra1-3xHA protein levels in whole cell lysates of the indicated *yra1* mutants at 30°C and 42°C. Pgk1 serves as loading control. (B and C) Quantification of total Yra1-3xHA protein levels for *yra1* single (B) and double mutants (C) at 30°C and 42°C, as shown in (A). Yra1 protein levels are normalized to Pgk1 and Yra1 wt is set to 1. Preventing Yra1 phosphorylation at both S8 and S142 modestly reduces Yra1 levels during heat stress, whereas the *yra1-S8A* single mutant has no such effect. Wilcoxon matched-pairs signed rank t-test was used to determine statistical significance: $^{*}p < 0.05$.

(D) Mex67 protein levels in total lysates are slightly increased in *yra1-S8A-S142A CBP20-TAP MEX67-9xmyc* cells after heat shock. Representative Western blots and quantification of lysates and Cbp20-bound nuclear mRNPs. Native purification of nuclear mRNPs via Cbp20-TAP comparing *YRA1-3xHA* cells with *yra1-S8D-S142D-3xHA* or *yra1-S8A-S142A-3xHA* cells at 30°C or 42°C, respectively. Total protein levels were normalized over protein levels in *CBP20-TAP YRA1-3xHA* cells. Cbp20-bound samples were normalized to the amount of purified Cbp20 and to the Cbp20-TAP *YRA1-3xHA* control for each replicate individually. Wilcoxon matched-pairs signed rank t-test was used to determine the statistical significance: $^{*}p < 0.05$, $^{**}p < 0.01$.

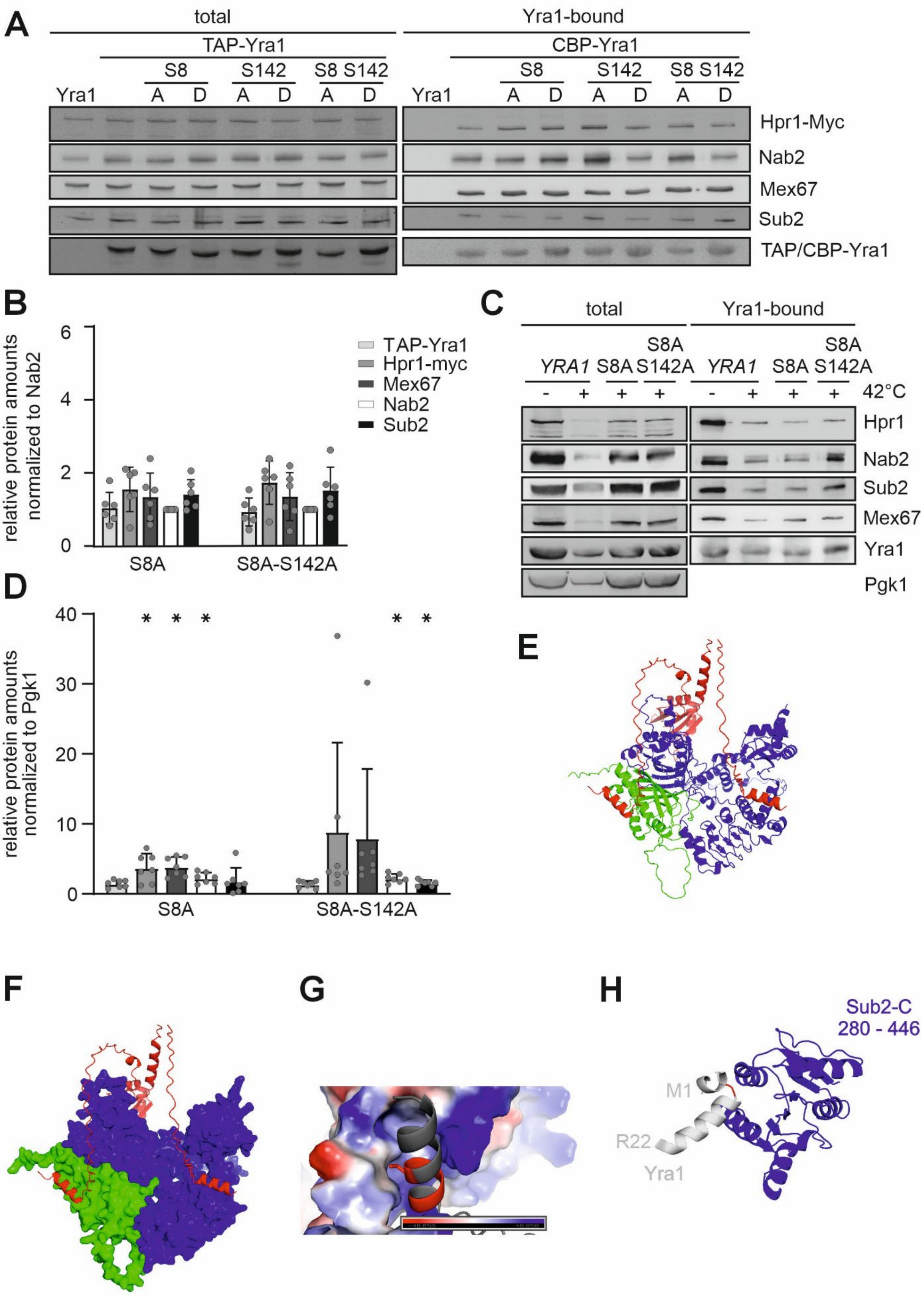
A
total
Yra1-bound
TAP-Yra1
CBP-Yra1
S8
S142
S8 S142
Yra1
A
D
Hpr1-Myc
Nab2
Mex67
Sub2
TAP/CBP-Yra1
B
relative protein amounts normalized to Nab2
TAP-Yra1
Hpr1-myc
Mex67
Nab2
Sub2
S8A
S8A-S142A
C
total
Yra1-bound
YRA1
S8A
S8A S142A
42°C
Hpr1
Nab2
Sub2
Mex67
Yra1
Pgk1
D
relative protein amounts normalized to Pgk1
S8A
S8A-S142A
E
F
G
H
Sub2-C
280 - 446
M1
R22
Yra1

**Figure S4**. **The interaction between Yra1 and Mex67 is increased in yra1-S8A *cells*, related to Figure 5**

(A) Representative Western blots of whole cell extracts (total) used for tandem affinity purification of different Yra1 phosphorylation site mutants grown at 30°C. For the quantification results of Yra1-bound samples see Figure 5 C.

(B) Quantification of protein levels shown in (A). Total protein levels were normalized to Nab2 and to *TAP-YRA1* control cells for each replicate. (C) Representative Western blots of total sample used for TAP-Yra1 purification after heat shock (42°C). For the quantification result of Yra1-bound samples see Figure 5 D.

(D) Quantification of proteins levels shown in (C). Total protein levels were normalized to Pgk1 and to *TAP-YRA1* control cells for each replicate. Wilcoxon matched-pairs signed rank t-test was used to determine the statistical significance: $^{*}p < 0.05$.

(E to G) The N-terminal Yra1 phosphorylation site S8 localizes within the Mex67 interacting motif. AlphaFold prediction (Abramson et al. 2024) of the interaction between Yra1 (red) with Mex67 (blue) and Mtr2 (green) (E and F) shows binding of Mex67 to the N-terminal UBM of Yra1. Binding surface of the Yra1 N-terminus (represented as ribbon diagram in gray and SLDD motif in red) with Mex67 (surface charge) (G).

(H) The N-terminal Yra1 phosphorylation site S8 interacts with Sub2-C. AlphaFold[1] prediction of the interaction between Yra1 (gray) with Sub2 (blue) shows binding of Sub2 to the N-terminal UBM of Yra1. Only residues 1-22 from Yra1 and 280-446 from Sub2 are shown. Yra1 S8 is highlighted in red.

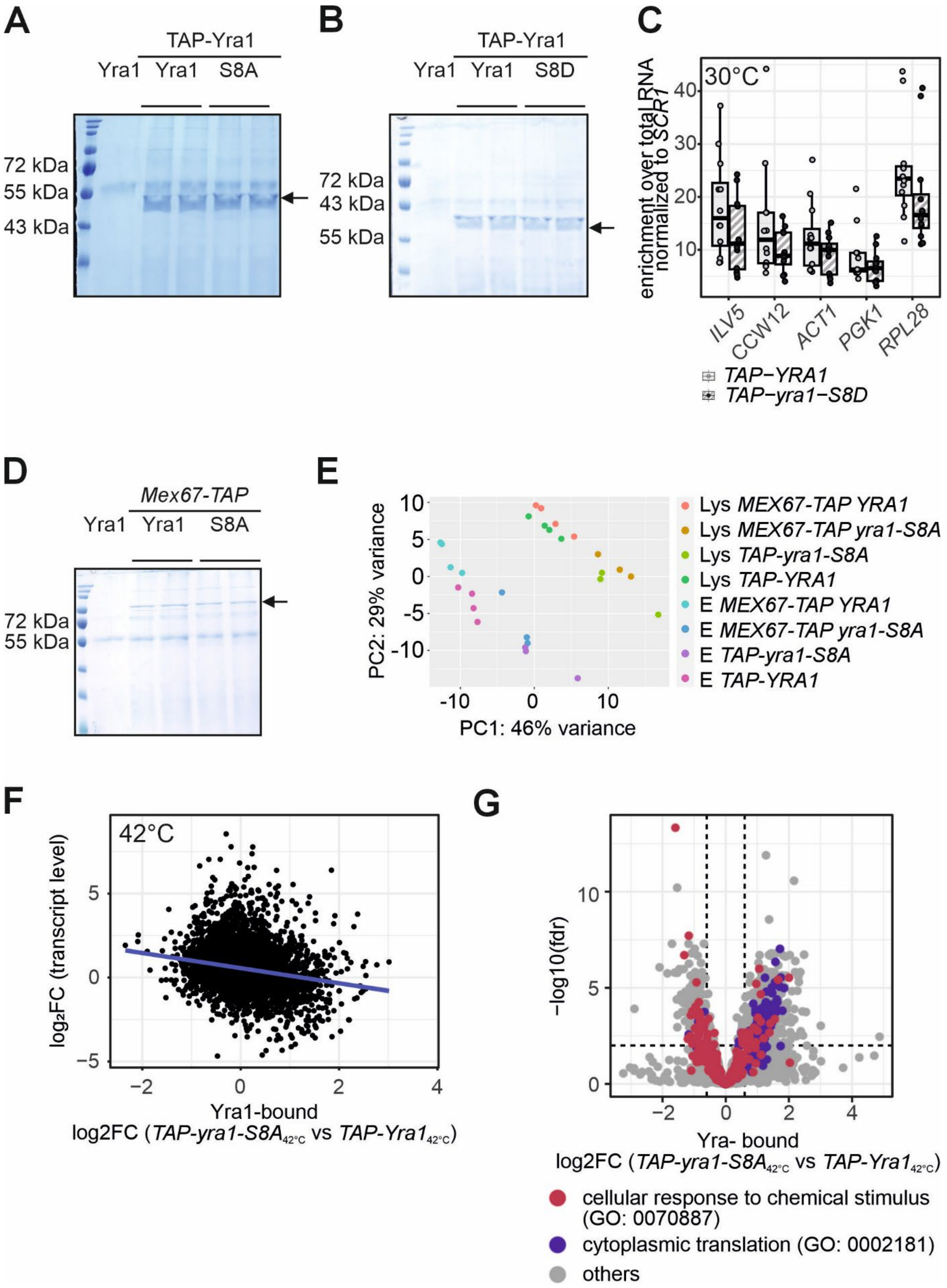
A
TAP-Yra1
Yra1 Yra1 S8A
72 kDa
55 kDa
43 kDa
B
TAP-Yra1
Yra1 Yra1 S8D
72 kDa
43 kDa
55 kDa
C
30°C
enrichment over total RNA normalized to SCR1
40
30
20
10
ILV5
CCW12
ACT1
PGK1
RPL28
TAP−YRA1
TAP−yra1−S8D
D
Mex67-TAP
Yra1 Yra1 S8A
72 kDa
55 kDa
E
PC2: 29% variance
PC1: 46% variance
10
5
0
-5
-10
-10
0
10
Lys MEX67-TAP YRA1
Lys MEX67-TAP yra1-S8A
Lys TAP-yra1-S8A
Lys TAP-YRA1
E MEX67-TAP YRA1
E MEX67-TAP yra1-S8A
E TAP-yra1-S8A
E TAP-YRA1
F
42°C
log₂FC (transcript level)
5
0
−5
−2
0
2
4
Yra1-bound
log2FC (TAP-yra1-S8A42°C vs TAP-Yra142°C)
G
−log10(fdr)
10
5
0
−2
0
2
4
Yra- bound
log2FC (TAP-yra1-S8A42°C vs TAP-Yra142°C)
cellular response to chemical stimulus (GO: 0070887)
cytoplasmic translation (GO: 0002181)
others

**Figure S5. Yra1 RNA binding is unchanged in *yra1-S8D* cells at 30°C, related to Figure 5**
(A) Yra1 amounts purified to determine the amount of bound RNA by RNA immunoprecipitation (RIP) are similar for *yra1-S8A* and *YRA1* cells. Representative Coomassie gels from RIP experiments to determine RNA binding by Yra1 grown at 42°C. The arrows indicate TAP-Yra1. For RT-qPCR results see Figure 5.
(B) Yra1 amounts purified to determine the amount of bound RNA by RNA immunoprecipitation (RIP) are similar for TAP-*yra1-S8D* and *TAP-YRA1* cells. Representative Coomassie gel showing TAP-Yra1 RIP eluates from cells grown at 30°C. The arrow indicates TAP-Yra1.
(C) RT-qPCR to determine the levels of specific transcripts bound to Yra1 of the experiment shown in (A). The enrichment was calculated by normalizing CT values of the respective mRNAs over the CT values of the ncRNA *SCR1* and to the respective lysate samples. $n \geq 6$, * $p < 0.05$.
(D) Mex67 amounts purified to determine the amount of bound RNA by RNA immunoprecipitation (RIP) are similar for *yra1-S8A* and *YRA1* cells. Representative Coomassie gels from RIP experiments to determine RNA binding by Mex67 grown at 42°C. The arrows indicate Mex67-TAP. For RT-qPCR results see Figure 5.
(E) Total RNA and Yra1- or Mex67-bound RNA separate in Principle component analysis (PCA) of the RIP sequencing data generated using Deseq2 from Galaxy. Lys: total RNA, E: Yra1- or Mex67-bound RNA. $n \geq 3$.
(F) Transcripts that are more abundant during heat stress are less efficiently bound by yra1-S8A. Correlation between the $\log_2$FC of TAP-yra1-S8A- vs. TAP-YRA1-bound to the $\log_2$FC of total transcript abundance after shift of the cells from 25°C to 37°C for each transcript contained in both data sets (total RNA amounts t16 vs. t0 taken from Castells-Roca et al. (2011))[3]. For analysis after normalization to the total RNA see Figure S5.
(G) Transcripts encoding proteins involved in cytoplasmic translation (blue; GO:0002181) exhibit increased binding to TAP-yra1-S8A, whereas transcripts associated with cellular responses to chemical stimuli (red; GO:0070887) show reduced binding. Volcano plot showing differential association of transcripts with TAP-yra1-S8A relative to TAP-Yra1.

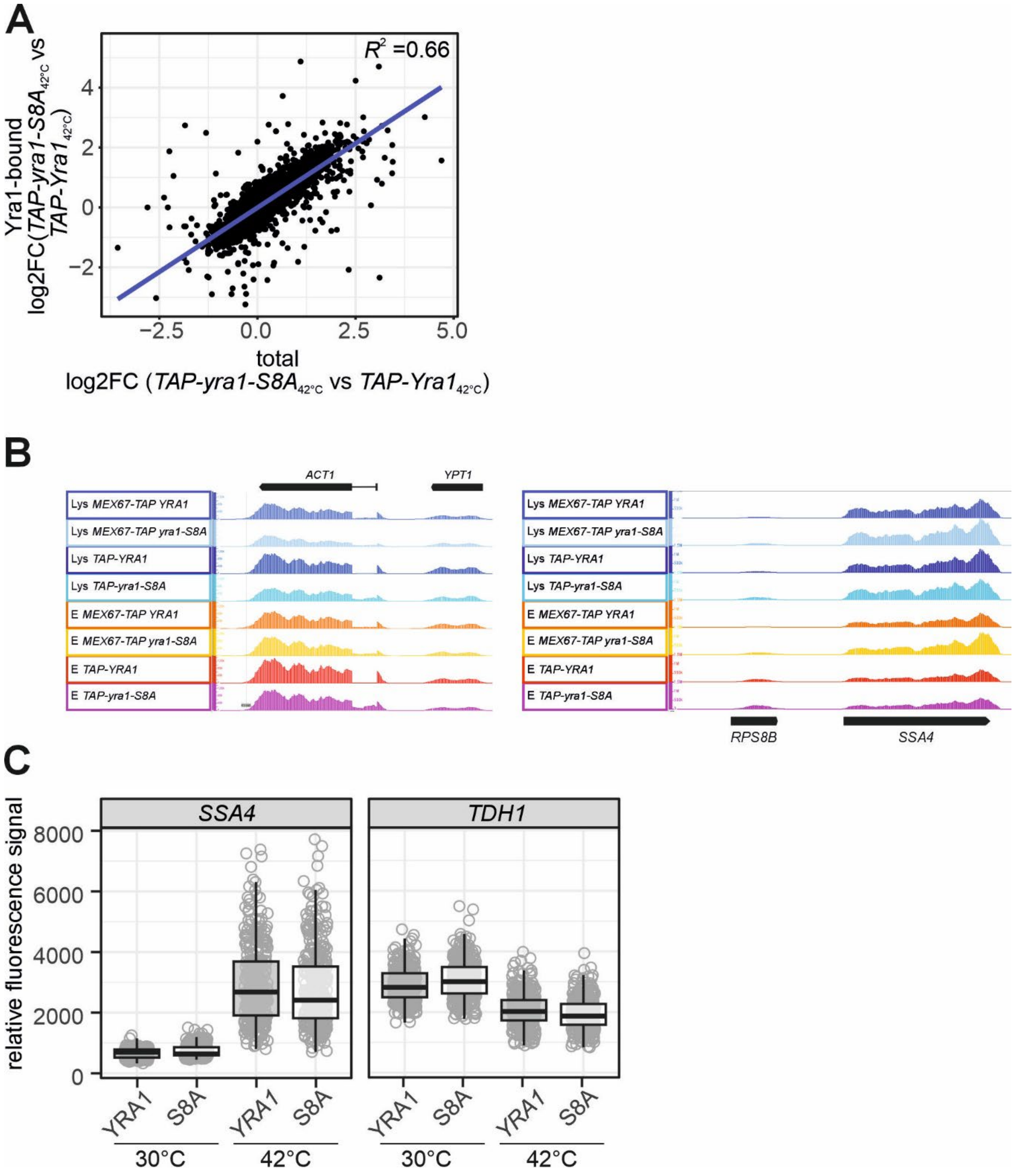

A
$R^2$ =0.66
Yra1-bound
log2FC(TAP-yra1-S8A42°C vs TAP-Yra142°C)
total
log2FC (TAP-yra1-S8A42°C vs TAP-Yra142°C)
B
ACT1
YPT1
Lys MEX67-TAP YRA1
Lys MEX67-TAP yra1-S8A
Lys TAP-YRA1
Lys TAP-yra1-S8A
E MEX67-TAP YRA1
E MEX67-TAP yra1-S8A
E TAP-YRA1
E TAP-yra1-S8A
RPS8B
SSA4
C
SSA4
TDH1
relative fluorescence signal
YRA1
S8A
30°C
42°C

**Figure S6. Intron-containing RNA accumulates in *yra1-S8A* cells, related to Figure 5 and 6**

(A) Changes in transcript binding by yra1-S8A are caused by changes in transcript abundance. Scatter plot displaying the correlation between the changes in the Yra1-bound and total RNA of *TAP-yra1-S8A* over *TAP-YRA1* cells.

(B) RNA from *yra1-S8A* cells has more reads mapping to introns, e.g. of the *ACT1* transcript, while Yra1- and Mex67-binding to the stress-induced transcript *SSA4* is not altered. Mapped sequencing reads of the intron-containing *ACT1* transcript and the heat shock-induced *SSA4* transcript for total RNA (Lys: lysate) and Yra1- or Mex67-bound RNA (E: eluate) of *yra1-S8A* and wt *YRA1* cells.

(C) The total fluorescence increases and decreases during heat stress for *SSA4* and *TDH1* mRNA probes, respectively. Quantification of the fluorescence signal of each cell normalized by the mean signal intensity of the whole experiment.

**Table S1. Conditions for different stress treatments tested for the occurrence of a nuclear mRNA export inhibition, related to Figure 1**

| Stress | treatment |
|---|---|
| Heat Shock | Add hot media to reach 42°C, incubate for 15 min or 1 h depending on the experiment |
| Hyperosmotic stress | 1.25 M NaCl in YPD for 1 h |
| Hypoosmotic stress | Shift cells from 20% YPD 1 M sorbitol to 20% YPD 0.1 M sorbitol for 5 min |
| Ethanol stress | 8% ethanol in YPD for 1 h |
| Oxidative stress | 2 mM arsenite for 1 h |
| Rapamycin | 2 µg/ml rapamycin in ethanol with 20% Tween-20 for 1 h |
| Unfolded protein response | 8 mM DTT in YPD for 1 h |
| Glucose starvation | SC media without glucose for 1 h |
| Nitrogen starvation | 2% glucose, 0.19% (w/v) yeast nitrogen base without amino acids and without ammonium sulphate for 1 h |

**Table S2. mRNA export adaptors are differentially phosphorylated under stress, related to Figure 2**

Fold changes and corrected p-values (fdr) for the phosphorylated sites identified in the phosphoproteome analysis. Sites with a significant 2-fold change as well as almost 2-fold change are highlighted in black and grey (bold), respectively.

| | | heat stress | | hyperosmotic stress | | *Δslt*30-wt30 | | *Δslt*42-wt42 | |
|---|---|---|---|---|---|---|---|---|---|
| protein | position | FC | fdr | FC | fdr | FC | fdr | FC | fdr |
| Hpr1 | 227 | 1.14 | 2.48E-01 | 1.09 | 0.4179 | 1.15 | 0.3951 | 1.04 | 0.8798 |
| | 234 | 0.97 | 7.99E-01 | 0.76 | 0.0524 | 1.01 | 0.9782 | 1.08 | 0.7662 |
| | 673 | **0.28** | 9.22E-06 | **0.49** | 0.0002 | 0.88 | 0.4065 | 1.01 | 0.9570 |
| | 675 | **0.49** | 1.81E-04 | 0.67 | 0.0083 | 0.91 | 0.6350 | 0.92 | 0.7232 |
| | 694 | 0.80 | 2.38E-02 | **1.89** | 3.39E-05 | 0.81 | 0.1200 | 1.00 | 0.9952 |
| Npl3 | 224 | 0.80 | 8.56E-03 | 0.75 | 0.0020 | 0.90 | 0.3422 | 0.99 | 0.9528 |
| | 227 | 0.92 | 3.33E-01 | 0.91 | 0.2822 | 0.99 | 0.9441 | 1.07 | 0.6478 |
| | 349 | **2.74** | 3.58E-04 | **0.47** | 0.0027 | 1.94 | 0.0379 | 1.12 | 0.7805 |
| Nab2 | 2 | 0.70 | 7.17E-04 | 0.77 | 0.0053 | 1.11 | 0.3571 | 1.12 | 0.3677 |
| | 254 | **2.03** | 3.42E-04 | **1.88** | 0.0009 | 1.00 | 0.9967 | 0.69 | 0.0746 |
| Yra1 | 8 | **1.97** | 6.58E-05 | 0.88 | 0.2661 | 0.92 | 0.6518 | **0.48** | 0.0003 |
| | 142 | **1.79** | 4.78E-04 | **2.89** | 7.06E-06 | 1.06 | 0.7776 | 0.78 | 0.1798 |
| | 149 | 0.71 | 5.16E-02 | **0.51** | 0.0017 | 0.83 | 0.4572 | 0.74 | 0.2237 |
| | 211 | 1.17 | 7.25E-02 | **7.06** | 5.90E-08 | 0.62 | 0.0034 | 0.92 | 0.5256 |

**Table S3. Plasmids used in this study**

| Plasmid | Description | Reference |
|---|---|---|
| pRS313 | | [2] |
| pRS314 | | [2] |
| pRS315 | | [2] |
| pRS316 | | [2] |
| pRS315-*YRA1* | Plasmid containing the Yra1 coding region with 300 bp up and downstream in a pRS315 plasmid | [4] |
| pRS316-*YRA1* | Plasmid containing the Yra1 coding region with 300 bp up and downstream in a pRS316 plasmid | [4] |

| | | |
|---|---|---|
| pRS315-*yra1-S8A* | Yra1 plasmid contains a mutation that prevents phosphorylation at position S8 of Yra1 | This study |
| pRS315-*yra1-S142A* | Yra1 plasmid contains a mutation that prevents phosphorylation at position S142 of Yra1 | This study |
| pRS315-*yra1-S211A* | Yra1 plasmid contains a mutation that prevents phosphorylation at position S211 of Yra1 | This study |
| pRS315-*yra1-S8A-S142A* | Yra1 plasmid contains a mutation that prevents phosphorylation at position S8 and S142 of Yra1 | This study |
| pRS315-*yra1-S8D-S142D* | Yra1 plasmid contains a phosphomimicry mutation at position S8 and S142 of Yra1 | This study |
| pRS315-TAP-*yra1* | Plasmid with N-terminal TAP-tagged Yra1 | This study |
| pRS315-TAP-*yra1-S8A* | Plasmid with N-terminal TAP-tagged Yra1 contains a mutation that prevents phosphorylation at position S8 of Yra1 | This study |
| pRS315-TAP-*yra1-S142A* | Plasmid with N-terminal TAP-tagged Yra1 contains a mutation that prevents phosphorylation at position S142 of Yra1 | This study |
| pRS315-TAP-*yra1-S8A-S142A* | Plasmid with N-terminal TAP-tagged Yra1 contains a mutation that prevents phosphorylation at position S8 and S142 of Yra1 | This study |
| pRS315-TAP-*yra1-S8D* | Plasmid with N-terminal TAP-tagged Yra1 contains a phosphomimicry mutation at position S8 of Yra1 | This study |
| pRS315-TAP-*yra1-S142D* | Plasmid with N-terminal TAP-tagged Yra1 contains a phosphomimicry mutation at position S142 of Yra1 | This study |
| pRS315-TAP-*yra1-S8D-S142D* | Plasmid with N-terminal TAP-tagged Yra1 contains a phosphomimicry mutation at position S8 and S142 of Yra1 | This study |
| pRS315-*yra1*-3xHA | Plasmid with C-terminal 3xHA-tagged Yra1 | This study |
| pRS315-*yra1-S8A*-3xHA | Plasmid with C-terminal 3xHA-tagged Yra1 contains a mutation that prevents phosphorylation at position S8 of Yra1 | This study |
| pRS315-*yra1-S142A*-3xHA | Plasmid with C-terminal 3xHA-tagged Yra1 contains a mutation that prevents phosphorylation at position S142 of Yra1 | This study |
| pRS315-*yra1-S211A*-3xHA | Plasmid with C-terminal 3xHA-tagged Yra1 contains a mutation that prevents phosphorylation at position S211 of Yra1 | This study |
| pRS315-*yra1-S8A-S142A*-3xHA | Plasmid with C-terminal 3xHA-tagged Yra1 contains a mutation that prevents phosphorylation at position S8 and S142 of Yra1 | This study |
| pRS315-*yra1-S8D*-3xHA | Plasmid with C-terminal 3xHA-tagged Yra1 contains a phosphomimicry mutation at position S8 of Yra1 | This study |
| pRS315-*yra1-S142D*-3xHA | Plasmid with C-terminal 3xHA-tagged Yra1 contains a phosphomimicry mutation at position S142 of Yra1 | This study |
| pRS315-*yra1-S211D*-3xHA | Plasmid with C-terminal 3xHA-tagged Yra1 contains a phosphomimicry mutation at position S211 of Yra1 | This study |
| pRS315-*yra1-S8D-S142D*-3xHA | Plasmid with C-terminal 3xHA-tagged Yra1 contains a phosphomimicry mutation at position S8 and S142 of Yra1 | This study |
| pRS315-*yra1-S142D-S211D*-3xHA | Plasmid with C-terminal 3xHA-tagged Yra1 contains a phosphomimicry mutation at position S142 and S211 of Yra1 | This study |
| pRS315-*yra1-S142A-S211A*-3xHA | Plasmid with C-terminal 3xHA-tagged Yra1 contains a mutation that prevents phosphorylation at position S142 and S211 of Yra1 | This study |
